# Yra2 regulates proteolysis of Cse4 to prevent its mislocalization to non-centromeric regions for chromosomal stability in budding yeast

**DOI:** 10.64898/2026.08.11.744167

**Authors:** Prashant K. Mishra, Kentaro Ohkuni, Pascal Raymond, Michael Costanzo, Charles Boone, Daniel R. Zenklusen, Munira A. Basrai

**Affiliations:** Yeast Genome Stability Section, Genetics Branch, NCI/NIH, Bethesda, MD, USA; University of Montreal, Montreal, QC, Canada; University of Toronto, Toronto, ON, Canada

**Author notes:** Corresponding author: Munira A. Basrai, Ph.D., Genetics Branch, National Cancer Institute, National Institutes of Health, 41 Medlars Drive Rm B900, Bethesda, MD, 20892 USA.

**Keywords:** Centromere, kinetochore, chromosomal instability, Cse4, CENP-A, RNA export, proteolysis

## Abstract

Restricting the localization of centromere-specific histone H3 variant Cse4 (CENP-A in humans) to centromeric chromatin is essential for chromosome segregation. Mislocalization of overexpressed Cse4/CENP-A to non-centromeric regions contributes to chromosomal instability (CIN) in model organisms and human cells. CIN is an important hallmark of many cancers and hence defining mechanisms that prevent mislocalization of Cse4 is clinically significant. Here we report a role for *YRA2* (<u>Y</u>east <u>R</u>NA <u>A</u>nnealing Protein 2) in ubiquitin mediated proteolysis of Cse4 to prevent its mislocalization for chromosomal stability. *YRA2* was identified in a genome-wide screen for gene deletions that exhibit synthetic dosage lethality (SDL) upon overexpression of *CSE4* (*GALCSE4*). We determined that *yra2Δ* strains exhibit increased Cse4 stability, enriched Cse4 chromatin association, reduced Cse4 ubiquitination, Cse4 mislocalization, and CIN. Defects in interaction of E3 ubiquitin ligase Psh1 with Cse4 contributes to stability of Cse4 in *yra2Δ* strains. Consistent with these results, overexpression of *PSH1* suppresses *GALCSE4* SDL in *yra2Δ* strain. We determined that Yra2 mediated proteolysis of Cse4 is independent of its RNA related functions as strain deleted for the C-terminal ChTOP domain of Yra2 with an intact N-terminal RNA binding domain exhibits *GALCSE4* SDL and defects in Cse4 proteolysis. Furthermore, poly(A)^+^ RNA export mutants in *YRA1* (*yra1-2*) and *MEX67* (*mex67-5),* that interact with Yra2, do not exhibit *GALCSE4 SDL* and defects in RNA export are not observed in *yra2Δ* cells. In summary, we have defined a key role for Yra2 in preventing mislocalization of Cse4 by facilitating its proteolysis to preserve chromosomal stability.

**Article summary:** Accurate segregation of chromosomes during cell division is essential because segregation errors are linked to cancer and developmental disorders. We investigated how cells prevent mislocalization of centromere-specific histone H3 variant Cse4, which is essential for faithful chromosome segregation. We found that the yeast RNA annealing protein Yra2 prevents Cse4 mislocalization by promoting Psh1 mediated ubiquitination and degradation of Cse4. Cells lacking Yra2 showed increased stability of Cse4, enhanced chromatin enrichment with mislocalization to non-centromeric regions and CIN. These defects were suppressed by induction of Psh1. Our findings reveal a novel role for Yra2 in regulating Cse4 levels for chromosomal stability.

## Introduction

Kinetochores (Centromeric DNA and associated proteins) are essential for faithful chromosome segregation during the cell division (KITAGAWA AND HIETER 2001; MERALDI *et al*. 2006; LAWRIMORE *et al*. 2011; BURRACK AND BERMAN 2012; MADDOX *et al*. 2012; BIGGINS 2013; SCOTT AND BLOOM 2014; BARRA AND FACHINETTI 2018; SALINAS-LUYPAERT AND FACHINETTI 2024; DUTTA *et al*. 2025; KOCH AND MARSTON 2025). Centromeric DNA sequences are typically highly repetitive and AT-rich across species, and are occupied by a single centromeric nucleosome in budding yeast or multiple centromeric nucleosomes in other eukaryotes (KITAGAWA AND HIETER 2001; MERALDI *et al*. 2006; VERDAASDONK AND BLOOM 2011; BURRACK AND BERMAN 2012; MADDOX *et al*. 2012; BIGGINS 2013; BLOOM AND COSTANZO 2017; FRIEDMAN AND FREITAG 2017; Barra and Fachinetti 2018; Sidhwani and Straight 2023; Salinas-Luypaert and FACHINETTI 2024; DUTTA *et al*. 2025; GRECO *et al*. 2026). These centromeric nucleosomes are defined epigenetically by a specific histone H3 variant that is evolutionarily conserved from yeast to humans (Cse4 in *Saccharomyces cerevisiae*, Cnp1 in *Schizosaccharomyces pombe*, CID in fruit flies, and CENP-A in humans) (MCKINLEY AND CHEESEMAN 2016; SRIVASTAVA *et al*. 2018; SHARMA *et al*. 2019). Notably, overexpressed CENP-A leads to its mislocalization to non-centromeric regions, which correlates with the increased chromosomal instability (CIN) in yeasts, fruit flies, human cells and xenograft mouse model (HEUN *et al*. 2006; AU *et al*. 2008; MISHRA *et al*. 2011; AU *et al*. 2013; LACOSTE *et al*. 2014; ATHWAL *et al*. 2015; SHRESTHA *et al*. 2017; AU *et al*. 2020; EISENSTATT *et al*. 2021; SHRESTHA *et al*. 2021; OHKUNI *et al*. 2022; SHRESTHA *et al*. 2023; BALACHANDRA *et al*. 2024; SETHI *et al*. 2024; ZHANG *et al*. 2024; BALACHANDRA *et al*. 2025; SETHI *et al*. 2025). Intriguingly, CENP-A overexpression and ectopic mislocalization are frequently observed across a range of human cancers and this correlates with poor prognosis and drug resistance (TOMONAGA *et al*. 2003; AMATO *et al*. 2009; LI *et al*. 2011; MCGOVERN *et al*. 2012; SUN *et al*. 2016; ZHANG *et al*. 2016; SAHA *et al*. 2020; XU *et al*. 2020). Given the clinical significance of CENP-A, defining the molecular mechanisms that prevent mislocalization of CENP-A is an active area of investigation.

Studies from budding yeast have uncovered a role for molecular pathways that regulate chromosomal localization of Cse4 (GRECO *et al*. 2026). Post-translational modifications (PTMs), including ubiquitination, sumoylation, phosphorylation, and proline isomerization regulate cellular levels of Cse4 to prevent its mislocalization for chromosomal stability (COLLINS *et al*. 2004; AU *et al*. 2008; HEWAWASAM *et al*. 2010; RANJITKAR *et al*. 2010; BOECKMANN *et al*. 2013; HEWAWASAM *et al*. 2014; OHKUNI *et al*. 2014; OHKUNI *et al*. 2016; OHKUNI *et al*. 2018; EISENSTATT *et al*. 2020; OHKUNI *et al*. 2020; OHKUNI *et al*. 2022). Our genome-wide Synthetic Genetic Array (SGA) screen for gene deletions/mutants that exhibit synthetic dosage lethality (SDL) upon Cse4 overexpression have identified E3 ubiquitin ligases Psh1, F-box proteins SCF-Met30 and SCF-Cdc4, histone chaperone Hir2, Dbf1-dependent kinase (DDK) and Cdc48 segregase in preventing mislocalization of Cse4 to non-centromeric regions (CIFTCI-YILMAZ *et al*. 2018; AU *et al*. 2020; EISENSTATT *et al*. 2020; OHKUNI *et al*. 2022). Psh1 plays a central role in the proteolysis of overexpressed Cse4 (HEWAWASAM *et al*. 2010; RANJITKAR *et al*. 2010; CIFTCI-YILMAZ *et al*. 2018; EISENSTATT *et al*. 2020; OHKUNI *et al*. 2022). This process requires interactions with several key factors such as Spt16 (component of the FACT complex), Casein kinase 2 (CKA2), histone chaperone Hir2 and DDK (DEYTER AND BIGGINS 2014; HEWAWASAM *et al*. 2014; HILDEBRAND AND BIGGINS 2016; CIFTCI-YILMAZ *et al*. 2018; EISENSTATT *et al*. 2020). In contrast, under endogenous condition, Cdc48 segregase (along with cofactors Ufd1 and Npl4) acts to remove Psh1 modified-polyubiquitinated Cse4 from non-centromeric chromatin to prevent stable association of Cse4 at these ectopic sites (OHKUNI *et al*. 2022). Mck1 mediated phosphorylation of Cse4 facilitates the interaction between Cse4 and the SCF-Cdc4 E3 ubiquitin ligase complex for Cse4 proteolysis (ZHANG *et al*. 2024). In addition to ubiquitination, SUMO ligases (Siz1/Siz2) and the SUMO-targeted ubiquitin ligase Slx5 regulate Cse4 proteolysis independently of Psh1 (OHKUNI *et al*. 2016; OHKUNI *et al*. 2018). Ubiquitin-mediated proteolysis prevents mislocalization of Cse4 homologs in fission yeast, fruit flies and humans, highlighting the evolutionary conservation of mechanisms regulating the cellular levels of Cse4 (ARISTIZABAL-CORRALES *et al*. 2019; MORENO-MORENO *et al*. 2019; SETHI *et al*. 2024).

In addition to PTMs, chromosomal localization of Cse4 is also regulated by specific chaperones, histone dosage and structural conformation of Cse4 (CAMAHORT *et al*. 2007; MIZUGUCHI *et al*. 2007; STOLER *et al*. 2007; DEYTER *et al*. 2017; HEWAWASAM *et al*. 2018; EISENSTATT *et al*. 2021; OHKUNI *et al*. 2024). For example, histone chaperones such as replication-independent histone chaperone (HIR) complex prevents mislocalization of Cse4 to non-centromeric regions and CIN phenotype (CIFTCI-YILMAZ *et al*. 2018), whereas chromatin assembly factor (CAF-1), a component of conserved, heterotrimeric histone chaperone complex, promotes Cse4 mislocalization (HEWAWASAM *et al*. 2018). Moreover, several studies have defined a role for dosage of histone H4 in Cse4 localization (DEYTER *et al*. 2017; EISENSTATT *et al*. 2021). We have shown that histone H4 binds to Cse4 and disrupts an intramolecular interaction of Cse4 dimers, shifting Cse4 from a "closed" to an "open" conformation (OHKUNI *et al*. 2024). This "open" state facilitates Cse4 sumoylation and mislocalization to non-centromeric chromatin leading to CIN phenotype (OHKUNI *et al*. 2024). Reduced gene dosage of histone H4 suppresses the mislocalization of Cse4 in *psh1Δ* strains, likely by minimizing this Cse4-H4 interaction (EISENSTATT *et al*. 2021). In contrast, increased histone H4 dosage facilitates this interaction, promoting an "open" conformational state of Cse4 that leads to enhanced sumoylation and subsequent mislocalization (EISENSTATT *et al*. 2021; OHKUNI *et al*. 2024). Since Cse4 is not fully stabilized even when multiple known degradation pathways are simultaneously disrupted, additional mechanisms likely exist to regulate its proteolysis.

Our SGA screen identified *YRA2* (*<u>Y</u>*east *<u>R</u>*NA *<u>A</u>*nnealing Protein 2) as a significant negative genetic interactor for *GALCSE4* (CIFTCI-YILMAZ *et al*. 2018). Yra2 is a non-essential member of the REF (RNA Export Factor) family of RNA-binding proteins (ZENKLUSEN *et al*. 2001; ZENKLUSEN *et al*. 2002). Yra2 along with Yra1 and Mex67 are components of the RNA export pathway that help transport messenger RNA from the nucleus to the cytoplasm (ZENKLUSEN *et al*. 2001; ZENKLUSEN *et al*. 2002; KASHYAP *et al*. 2005). Yra1 is an essential RNA-binding export factor that plays a major role in gene transcription, RNA processing, and nuclear export (PORTMAN *et al*. 1997; ZENKLUSEN *et al*. 2002). Yra1 recruits Mex67–Mtr2 protein complex to processed mRNA transcripts that guide the transport of messenger ribonucleoprotein particles through nuclear pore complexes into the cytoplasm (ZENKLUSEN *et al*. 2001). Yra2 is a nonessential paralog of Yra1 (ZENKLUSEN *et al*. 2001). Although, Yra2 cannot fully replace the functions of Yra1, overexpression of *YRA2* can suppress the temperature sensitivity to growth phenotype of *yra1-2* cells, indicating partial functional redundancy between these two proteins (ZENKLUSEN *et al*. 2001; KASHYAP *et al*. 2005). Recent studies indicate that Yra2 has limited transcript occupancy and may function primarily as an adaptor or scaffold protein rather than as a major component of the RNA export machinery (ASADA *et al*. 2023). In addition, Yra2 is expressed at significantly lower levels than Yra1 (ZENKLUSEN *et al*. 2001), suggesting that Yra2 may possess specialized or additional functions in budding yeast.

In this study, we show a novel role for *YRA2* (<u>Y</u>east <u>R</u>NA <u>A</u>nnealing Protein 2) in ubiquitin mediated proteolysis of Cse4 to prevent its mislocalization for faithful chromosome segregation. Strain deleted of *YRA2* displays SDL upon overexpression of *CSE4* (*GALCSE4*). Mechanistically, this is characterized by impaired Cse4 ubiquitination that drives its stabilization and enrichment in the chromatin fraction, resulting in extensive non-centromeric mislocalization and subsequent CIN. Notably, Yra2-mediated proteolysis of Cse4 is independent of its RNA-related functions.

## Materials and methods

### Yeast strains, plasmids, and growth conditions

Yeast strains and plasmids used in this study are summarized in Table 1. Strains were cultured in YPD medium (1% yeast extract, 2% Bacto-peptone, 2% glucose) or in synthetic yeast medium supplemented with either 2% glucose or galactose+raffinose (2% each), along with the appropriate supplement for plasmid selection. Viability of wild-type and mutant strains carrying *GALCSE4* (pMB1458) or the control vector (pMB433 *GAL1*) and other plasmids was assessed by spotting serial dilutions of equal cell numbers from three transformants onto selective media and incubating at 25°C.

**Table 1.**
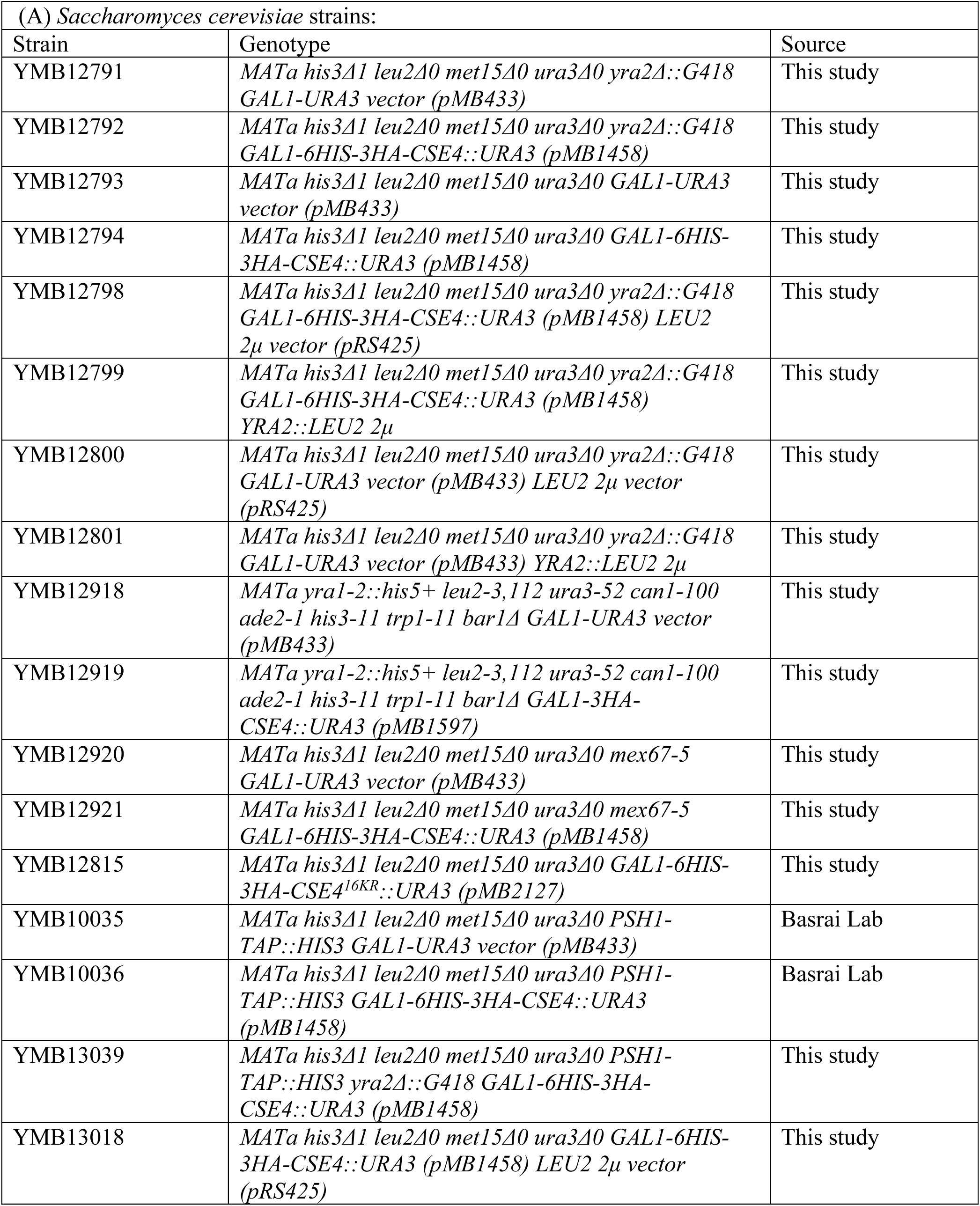
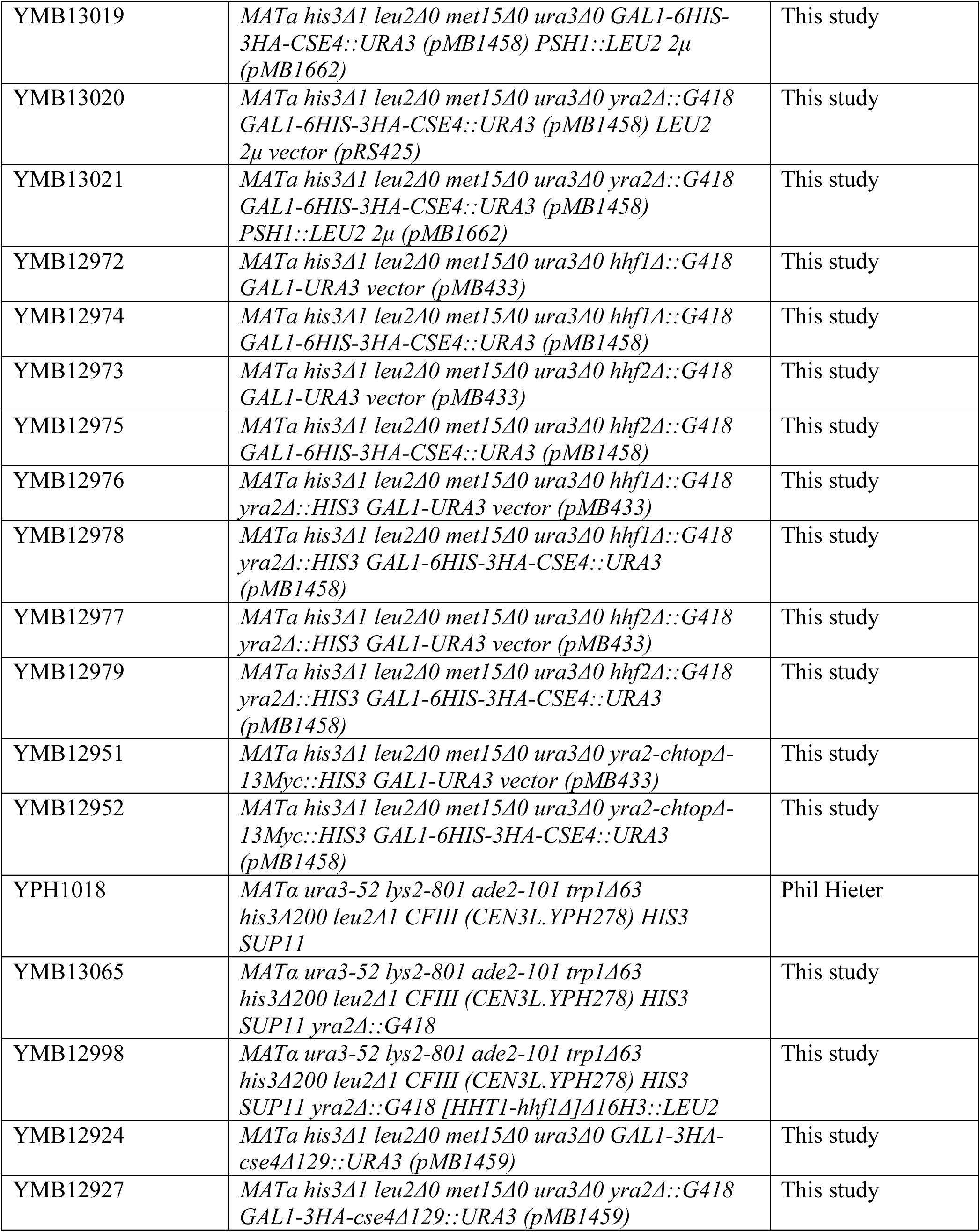
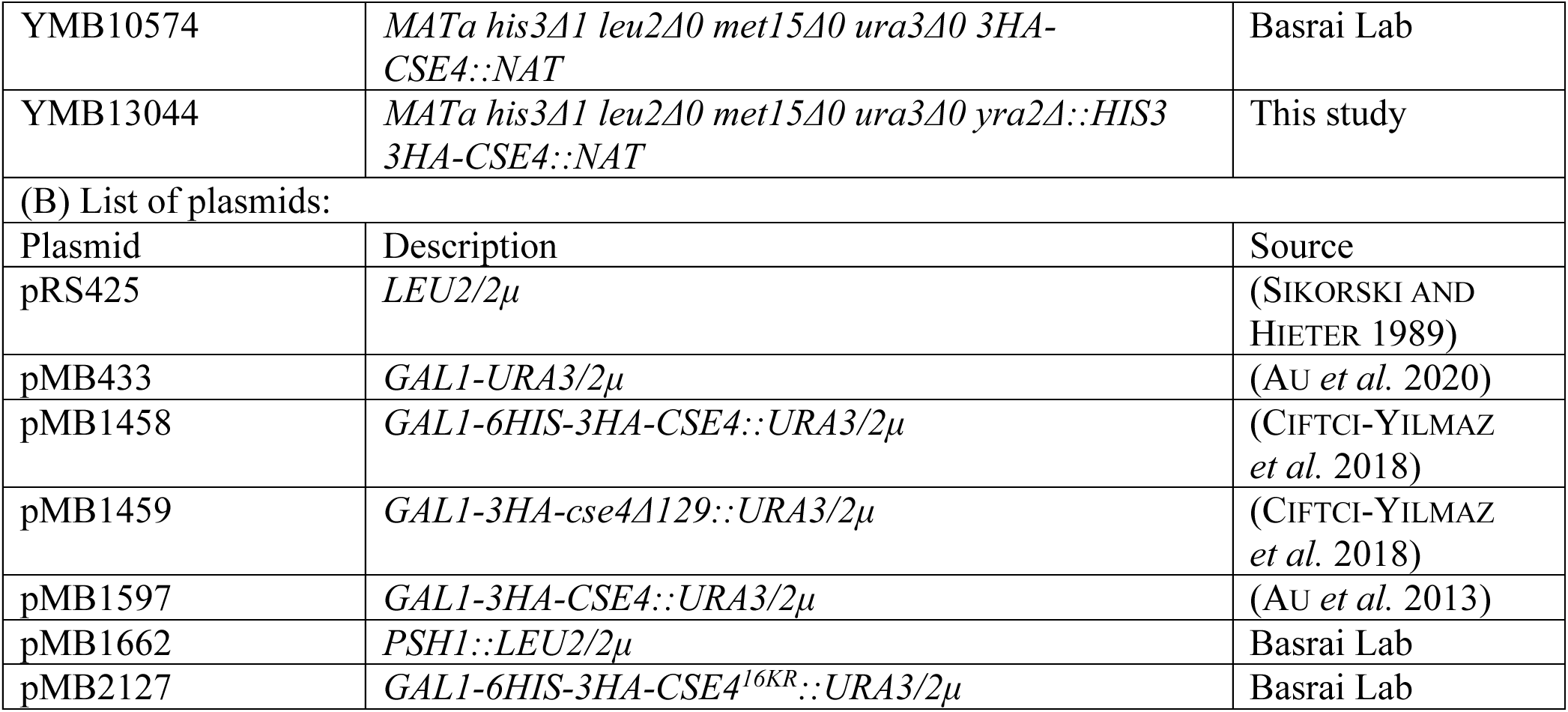
List of strains, and plasmids used in this study.

### Protein stability assays

Protein stability assays were done as previously described (AU *et al*. 2013; MISHRA *et al*. 2015; CIFTCI-YILMAZ *et al*. 2018). Wild-type and mutants carrying *HA-CSE4* expressed from *GAL1* promoter (*GALCSE4*, pMB1458) were grown to logarithmic phase at 25°C in selective media with 2% raffinose, added 2% galactose (2%) and grown for 3 hours to induce the expression of Cse4. Cells were collected at different time points after addition of 2% glucose and cycloheximide (CHX, 50 µg/mL) to block the protein translation. Protein extracts were prepared as described previously (KASTENMAYER *et al*. 2006), and equal amounts of proteins for each sample were prepared based on OD_750_ values determined by the Bio-Rad DC protein assay (500-0113, Bio-Rad Inc.) were size separated on a 4-12% Bis-Tris gel (Invitrogen Inc.) for western blot analysis. For protein stability of Cse4 expressed from its endogenous promoter, wild-type and *yra2*Δ strains were grown to logarithmic phase at 25°C in YPD and CHX (50 µg/mL) was added. Cells were collected at different time points, proteins extracts were prepared, equal amount determined by OD_750_ measurements, and analyzed by western blotting as described above. Primary antibodies included anti-HA (12CA5, Roche) to detect HA-tagged Cse4, and Tub2 (loading control), whereas secondary antibodies were HRP-conjugated sheep α-mouse IgG (NA931V, Amersham Biosciences) and HRP-conjugated donkey α-rabbit IgG (NA934V, Amersham Biosciences). Western blot band intensities were quantified using Image J software (SCHNEIDER *et al*. 2012). Protein stability was expressed as the percentage of protein remaining (normalized to Tub2) at the indicated time points following CHX treatment, with the initial protein level set to 100%.

### ChIP and qPCR experiments

ChIP experiments were performed following the methodologies described previously (MISHRA *et al*. 2007; MISHRA *et al*. 2011; MISHRA *et al*. 2013). Briefly, cells were cross-linked with 1% formaldehyde for 15 min at room temperature and quenched with 125 mM glycine for 5 min. Pellets were collected by centrifugation, washed with TBS (150 mM NaCl, 20 mM Tris-HCl, pH 7.6), and converted to spheroplasts using Zymolyase 100T. Spheroplasts were washed once in post-spheroplasting buffer (1.2 M sorbitol, 20 mM Na-PIPES, pH 6.8, 1 mM MgCl₂) and three times in FA buffer (50 mM Na-HEPES, pH 7.6, 1 mM EDTA, 150 mM NaCl, 0.1% sodium deoxycholate, 1% Triton X-100) supplemented with protease inhibitors (P8215, Sigma). Samples were resuspended in FA buffer and sonicated on ice (30% amplitude; five 12 sec pulses, 2 min intervals) to yield DNA fragments averaging ∼400 bp. Soluble chromatin was subjected to immunoprecipitation using anti-HA agarose (A2095, Sigma), and Histone H3 (Ab1791, Abcam) antibodies. ChIP DNA was analyzed by qPCR using a 7500 Fast Real-Time PCR System (Applied Biosystems) with SYBR green detection using primers as described (OHKUNI *et al*. 2020; EISENSTATT *et al*. 2021). Enrichment was calculated as % input using the _ΔΔ_C_T_ method (LIVAK AND SCHMITTGEN 2001).

### Ubiquitin affinity pull down experiments

Ubiquitin (Ub) pull-down assay was performed to determine the levels of ubiquitinated Cse4 as described previously (AU *et al*. 2013; MISHRA *et al*. 2015). Briefly, cell pellets from logarithmically growing cultures of wild-type and *yra2Δ* strains with *GALCSE4* (pMB1458, induced for 3 hours) as well as control strains (untagged and *GALcse4^16KR^*) were collected and resuspended in lysis buffer (20 mM Na₂HPO₄, 20 mM NaH₂PO₄, 50 mM NaF, 10 mM β-glycerophosphate, 5 mM tetrasodium pyrophosphate, 5 mM N-ethylmaleimide, 2 mM EDTA, 1 mM DTT, 1 mM PMSF, 1% NP-40, and protease inhibitor cocktail [P8215, Sigma]). Whole cell lysates were prepared by bead beating cells using an equal volume of glass beads (425–600 μm) in a FastPrep-24 homogenizer (MP Biomedical) until 90% cell lysis were achieved. Equal amounts of lysates from wild-type, mutants and control strains were prepared based on OD_750_ measurements using the Bio-Rad DC protein assay (500-0113, Bio-Rad Inc.). A part of cell lysate for each sample was used as input and remaining was incubated overnight with agarose beads conjugated with tandem ubiquitin-binding entities (UM401 Agarose-TUBE1, Life Sensors Inc) on a roller at 4°C. Beads were washed three times (5 min each wash) with 1xTBST (Tris-buffered saline with 0.1% Tween-20) at room temperature and incubated at 100°C for 10 minutes in 2x Laemmli buffer to elute the bound proteins. The eluted proteins were size separated by western blotting on a 4–12% Bis-Tris gel and ubiquitinated Cse4 was visualized using an anti-HA antibody (12CA5, Roche). Western blots were analyzed to determine the protein signal intensities using Image J (SCHNEIDER *et al*. 2012) and these values were used to calculate the relative ubiquitination of Cse4 (ubiquitinated Cse4/input Cse4). Student’s t-test was used to determine the statistical significance of relative ubiquitination of Cse4 between the strains.

### Immunoprecipitation (IP) experiments

IP experiments were performed following the procedure as described previously (MISHRA *et al*. 2016; OHKUNI *et al*. 2024; MISHRA *et al*. 2025). Strains were grown overnight in selective medium containing 2% raffinose to logarithmic phase, cells were collected and inoculated into selective medium containing raffinose+galactose (2% each) and incubated at 25°C for 3 hours for experiments with *GALCSE4*. Cells were dissolved in extraction buffer (40mM Hepes, pH7.5, 350mM NaCl, 0.1% Tween, 10% glycerol, 1mM DTT, 1mM PMSF, and protease inhibitors [P8215, Sigma]) and whole cell extracts were prepared by bead beating using a FastPrep-24 homogenizer (MP Biomedical). An equal concentration of protein extracts based on OD_750_ measurements was incubated at 4°C overnight with antibodies dependent on experiments: anti-HA agarose (A2095, Sigma) for HA-Cse4, and IgG agarose (A2909, Sigma) for Psh1-TAP. The immunoprecipitated proteins were washed in 1xTBST three times (5 min each wash) and were eluted in 2x Laemmli buffer. Western blotting was performed using primary antibodies: anti-TAP (CAB1001, Invitrogen), anti-HA (H6906, Sigma and 12CA5, Roche), anti-histone H4 (ab10158, Abcam) and Tub2. Secondary antibodies were HRP-conjugated donkey α-rabbit IgG (NA934V, Amersham Biosciences) and sheep α-mouse IgG (NA931V, Amersham Biosciences).

### Chromatin fractionation experiments

Subcellular fractionation was performed to examine the levels of Cse4 in whole cell extracts, soluble and chromatin fractions following the procedure as described previously (AU *et al*. 2008). Wild-type and *yra2Δ* strains carrying *GALCSE4* (pMB1458) were grown at 25°C in selective media with 2% raffinose to logarithmic phase, added 2% galactose to induce the expression of Cse4 and grown for 3 hours. Proteins were analyzed by western blotting using anti-HA (12CA5, Roche), H2B (ab1790, Abcam) and Tub2 antibodies. Band signal intensities were determined using Image J (SCHNEIDER *et al*. 2012) and chromatin bound Cse4 was calculated for wild-type and *yra2Δ* strains after normalization to the signal intensities from H2B. Statistical significance was determined using the Student’s t-test.

### Chromosome Segregation Assays

The loss of a non-essential reporter chromosome (RC) was used as a readout to measure the fidelity of chromosome segregation using a colony color assay as described previously (SPENCER *et al*. 1990). In this assay, appearance of red sectors in an otherwise a white yeast colony represents the loss of the RC. Strains were grown in synthetic media to maintain the RC to the logarithmic phase; cells were collected, dilutions were prepared based on OD_600_ measurements, and the equal number of cells for each strain were plated on synthetic medium with limiting adenine at 25°C. The frequency of chromosome loss was measured by counting the number of colonies that were at least half red (represents the loss of the RC at the first cell division) and statistical significance was determined by the Student’s t-test.

### Poly(A)^+^ RNA fluorescence in situ hybridization (FISH)

Forty-five-nucleotide-long thymine DNA oligonucleotide probes modified with a 3′ amine group (dT45-3′ modification, Amino Linker C7; sequence: TTT TTT TTT TTT TTT TTT TTT TTT TTT TTT TTT TTT TTT TTT TTT-NH₂, Invitrogen) and coupled to Cy3 (PA13101, Cytiva, Amersham Cy3 NHS Ester, 1 mg) were used. 50 μg of dried modified-oligo(dT) were mixed with 100 μg of Cy3 dye, resuspended in labeling buffer (0.1M sodium bicarbonate, pH 9.0), and incubated overnight at room temperature in the dark. Cy3-labelled probes were purified using Qiagen QIAquick Nucleotide Removal columns (#28304, Qiagen) and stored at -20°C in the dark.

FISH was essentially performed as described (CASTELNUOVO *et al*. 2013). Briefly, liquid cultures were inoculated from freshly picked colonies grown on SD-URA glucose plates at 25°C and grown in 50 mL SD-URA medium containing 2% glucose at 25°C until reaching mid-log phase (OD₆₀₀ = 0.6-0.8). Cultures were then fixed by adding paraformaldehyde (#15714, EMS) to a final concentration of 4% for 45 minutes at room temperature with shaking. Following fixation, cells were washed three times with 10 mL Buffer B (1.2 M sorbitol, 100 mM KHPO₄, pH 7.5) and stored overnight at 4°C in the same buffer. After pelleting and removing the supernatant, cell walls were digested using 300 U of lyticase (in 1x PBS at 25,000 U/mL and stored at -20°C; #L2524, Sigma-Aldrich) until approximately 50% of cells turned dark under a phase contract microscope (approximately 10 minutes). Digested cells were dropped onto poly-L-lysine–coated 18 mm coverslips and allowed to settle overnight, washed once with Buffer B, and stored in 70% ethanol at -20 °C in 12-well plates. Before hybridization, ethanol was removed, cells were washed once with 2× SSC and equilibrated in 10% formamide/2× SSC. Probe hybridization mix containing 20 ng of dT-Cy3 probes was prepared in a solution of 10% formamide, 10% dextran sulfate, 2× SSC, 5 mM NaHPO₄ (pH 7.5), 0.5 mg/mL *E. coli* tRNA, and 0.5 mg/mL single-stranded DNA in a final volume of 20 μl per coverslip. Hybridization was carried out for 2 hours at 37°C, cells were washed twice in 10% formamide/2x SSC at 37 °C for 30 minutes, followed by brief DNA Hoechst staining (1x PBS containing 1 μg/mL Hoechst 33342, 2 min). Coverslips were dehydrated in 100% ethanol, dried, and mounted onto glass slides using ProLong Glass Antifade Mountant (#P36980, Invitrogen). Images were acquired using a Zeiss Axio Imager Z2 epifluorescence microscope equipped with a 100X/1.40 NA Plan-Apochromat oil-immersion objective, a Lumen Dynamics X-Cite 200 W metal-halide light source, and a 16-bit Photometrics Prime sCMOS camera. The following filter sets were used: Zeiss Set 49 for DAPI, Chroma SP102v1 for Cy3, and HR-DIC III optics. Multichannel z-stacks (240nm) were acquired and processed by deconvolution using default parameters in ZEN Blue (version 2.6 PRO, Zeiss). For visualization and analysis, 3D datasets were reduced to 2D using “Max Projection” in Image J. Cells were counted using Image J’s “Find Maxima” function with an appropriate threshold applied to the Hoechst channel and nuclear poly(A) accumulations scored manually as an increase in poly(A) signal co-localizing with Hoechst staining.

## Results

### Deletion of Yra2 exhibits SDL with overexpression of *CSE4*

We previously performed an SGA screen to identify gene deletions and mutants exhibiting SDL with *GALCSE4*, revealing roles for histone chaperones, E3 and SCF ubiquitin ligases, molecular segregases, and protein kinases in Cse4 proteolysis (CIFTCI-YILMAZ *et al*. 2018; AU *et al*. 2020; EISENSTATT *et al*. 2020; OHKUNI *et al*. 2020; EISENSTATT *et al*. 2021; OHKUNI *et al*. 2022; ZHANG *et al*. 2024) . While our previous genome-wide SGA screen successfully identified multiple protein classes required for Cse4 proteolysis, here we focused on *yra2Δ*, which was among the top candidates identified as a negative genetic interactor with *GALCSE4* (SGA score of -0.625 and *p* value of 8.70e-13) (CIFTCI-YILMAZ *et al*. 2018). To validate the results of the SGA screen, we transformed *GALCSE4* or empty vector into wild-type and *yra2*Δ strains and examined growth of these strains on glucose (*GALCSE4* OFF) or galactose (*GALCSE4* ON) medium. As expected, no growth defects were observed on galactose for strains transformed with vector alone. Consistent with the results of the screen, *yra2*Δ strain exhibit SDL with *GALCSE4* on galactose medium (Fig. 1a). We determined that the SDL of *GALCSE4* was linked to *yra2Δ*, as the growth defect of the *yra2Δ GALCSE4* strain on galactose medium was suppressed by expression of *YRA2* on a plasmid (Fig. 1b).

**Fig. 1.**
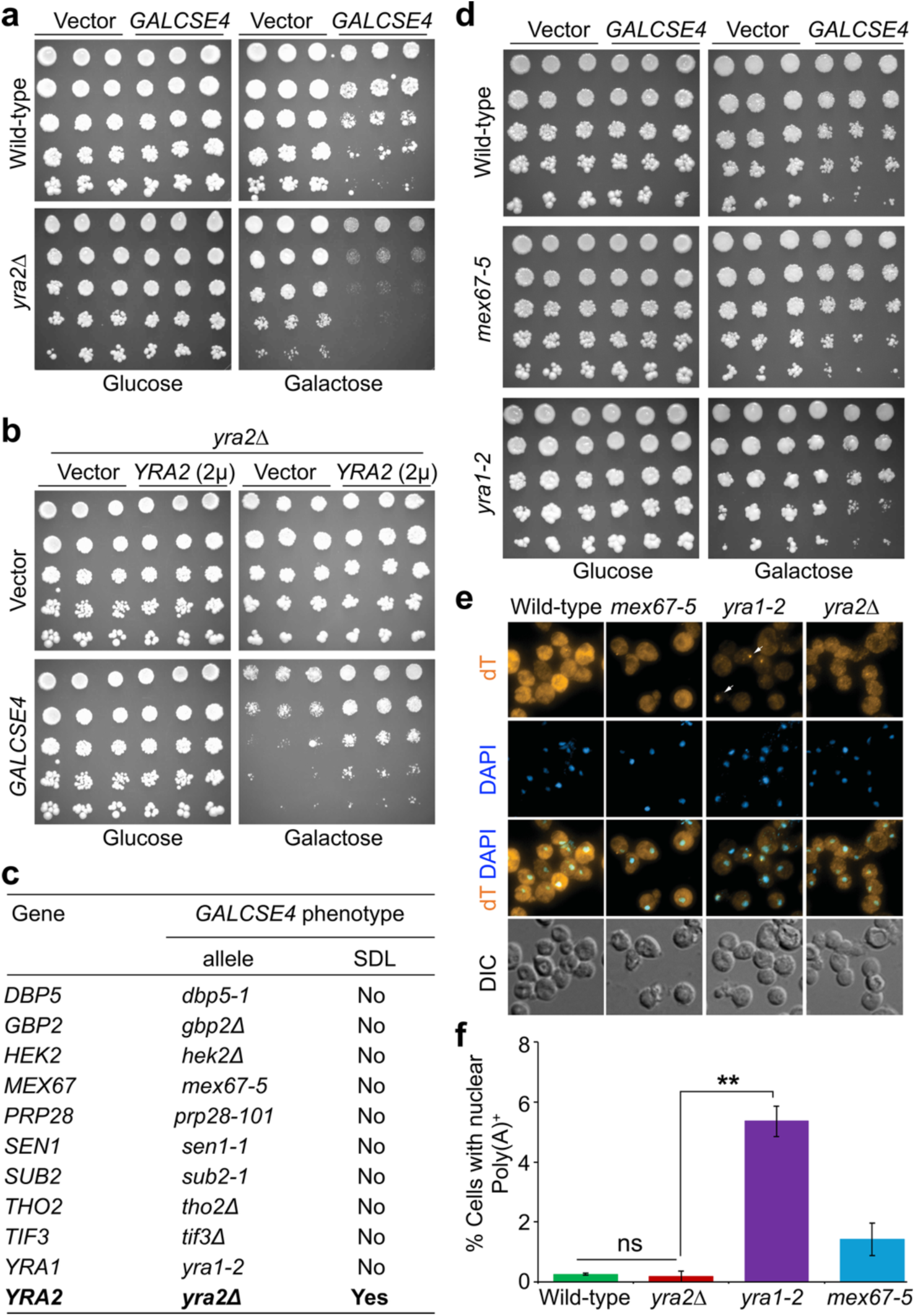
The loss of *YRA2* causes growth defects in strains overexpressing Cse4. a) Overexpression of Cse4 (*GALCSE4*) causes SDL in *yra2Δ* strains. Wild-type with vector [pMB433] (YMB12793) or *GALCSE4* [pMB1458] (YMB12794) and the isogenic *yra2Δ* with vector (YMB12791) or *GALCSE4* (YMB12792) were grown to logarithmic phase, five-fold serial dilutions were prepared and plated on SC-Ura plates containing either glucose (2%) or galactose+raffinose (2% each) at 25°C. b) Ectopic expression of *YRA2* suppresses the SDL phenotype in *yra2Δ GALCSE4* strains. *yra2Δ* strain with vectors [pMB433 and pRS425] (YMB12800); *GALCSE4* [pMB1458] + vector (YMB12798), vector + *YRA2/2μ* (YMB12801); and *GALCSE4* + *YRA2*/*2μ* (YMB12799) were grown to logarithmic phase, five-fold serial dilutions were prepared and plated on SC-Ura-Leu plates containing either glucose (2%) or galactose+raffinose (2% each) at 25°C. c) Mutants defective in RNA export does not exhibit SDL phenotype with *GALCSE4*. Results derived from previously published SGA screens with *GALCSE4* showed no significant SDL in mutants defective in RNA export (CIFTCI-YILMAZ *et al*. 2018; AU *et al*. 2020). d) *GALCSE4* does not exhibit SDL in *yra1-2* and *mex67-5* strains. Wild-type with vector (YMB12793) or *GALCSE4* (YMB12794), *yra1-2* with vector (YMB12918) or *GALCSE4* (YMB12919) and *mex67-5* with vector (YMB12920) or *GALCSE4* (YMB12921) were grown to logarithmic phase, five-fold serial dilutions were prepared and plated on SC-Ura plates containing either glucose (2%) or galactose+raffinose (2% each) at 25°C. e) Deletion of *YRA2* does not exhibit nuclear accumulation of poly(A)^+^ RNA. Wild-type (YMB12793), *yra1-2* (YMB12918), *mex67-5* (YMB12920) and *yra2Δ* (YMB12791) strains were grown as described in *Materials and metho*ds for the determination of poly(A)^+^ RNA. Representative images with arrow marking cells with a nuclear accumulation of poly(A)^+^ RNA are shown. f) Cells with nuclear poly(A)^+^ RNA in wild-type and mutant strains. Quantification were performed on three fields for each strain using strains in (e) and at least 300 cells were examined for each strain. Average±SE is shown, ns = statistically not significant, \*\**p* value <0.01, Student’s t-test.

Yra2 interacts *in vivo* with mRNA export factors Yra1 and Mex67 and overexpression of Yra2 suppresses the temperature sensitivity of *yra1-2* mutant strain (STRASSER AND HURT 2000; ZENKLUSEN *et al*. 2001; KASHYAP *et al*. 2005; OEFFINGER *et al*. 2007). We determined that deletion of *YRA1*, *MEX67*, and other known Yra2 interactors do not show growth defects with *GALCSE4* in our SGA screen (Fig. 1c). We further confirmed that *yra1-2* and *mex67-5* strains do not exhibit SDL phenotype with *GALCSE4* on galactose medium (Fig. 1d). No growth defects were observed for these strains transformed with vector alone on galactose plates (Fig. 1d). These results suggest that mutants impaired for mRNA export do not exhibit *GALCSE4* SDL, suggesting a previously unrecognized role for Yra2 in regulation of Cse4 in budding yeast.

### Yra2 is not required for export of poly(A)^+^ RNA from the nucleus

To further distinguish the role of Yra2 from those described for Yra1 and Mex67, we examined whether deletion of *YRA2* exhibits defect in the export of poly(A)^+^ RNA from the nucleus as reported for mutants of *YRA1* and *MEX67* (STRASSER AND HURT 2000; ZENKLUSEN *et al*. 2001). This was done by quantitative analysis of nuclear accumulation of poly(A)^+^ RNA in wild-type, *yra2Δ, yra1-2,* and *mex67-5* strains (ZENKLUSEN *et al*. 2001). As described previously (STRASSER AND HURT 2000; ZENKLUSEN *et al*. 2001), a significant nuclear accumulation of poly(A)^+^ RNA was observed in *yra1-2* and *mex67-5* mutants even at 25°C when compared to the wild-type strain (Fig. 1e and f). Quantitative analysis showed that for a wild-type strain, 0.25% of cells showed nuclear poly(A)^+^ RNA foci (0.25±0.029, average±SE), and this was significantly higher in *yra1-2* (5.36±0.511) and *mex67-5* (1.41±0.543) strains (Fig. 1f). However, no significant accumulation of nuclear poly(A)^+^ RNA was observed in *yra2Δ* strain (0.05±0.053) when compared to the wild-type strain (Fig. 1f). Taken together, these results show that the deletion of Yra2 does not impair the nuclear export of poly(A)^+^ RNA.

### Yra2 regulates ubiquitin-mediated proteolysis of Cse4

Previous studies have tightly correlated the *GALCSE4* SDL phenotype with defects in ubiquitin-mediated proteolysis, which consequently lead to increased stability of Cse4 and its subsequent enrichment in the chromatin fraction (AU *et al*. 2013; CIFTCI-YILMAZ *et al*. 2018; AU *et al*. 2020; EISENSTATT *et al*. 2020; ZHANG *et al*. 2024). Protein stability of overexpressed Cse4 in *yra2Δ GALCSE4* strain was examined after 3 hours of induction in galactose medium followed by addition of glucose and the protein synthesis inhibitor cycloheximide (CHX). Western blot analysis of whole cell extracts prepared at different time points after CHX treatment showed that Cse4 was rapidly degraded in the wild-type strain, however, the stability of Cse4 was significantly higher in the *yra2Δ* strain (∼2-fold, *p* value = <0.01) (Fig. 2a and b). To determine if the increased stability of Cse4 in whole cell extracts of *yra2Δ* strain is due to higher levels of Cse4 in the chromatin fraction, we performed subcellular fractionation with wild-type and *yra2Δ* strains. Consistent with previous studies with mutants defective for Cse4 proteolysis (CIFTCI-YILMAZ *et al*. 2018; AU *et al*. 2020), Cse4 was barely detectable in the soluble fraction of wild-type and *yra2Δ* strains (Fig. 2c). Our results showed that chromatin associated Cse4 was more enriched in the *yra2Δ* strain when compared to the wild-type strain (Fig. 2c and d). We have previously shown that the N-terminus of Cse4 is required for Cse4 proteolysis (AU *et al*. 2013), hence we asked if the N-terminus of Cse4 is required for the SDL of *GALCSE4*. Growth assays showed that *yra2*Δ strain transformed with *GALcse4Δ129* (deletion of the N-terminal 129 amino acids) do not exhibit SDL phenotype when compared to vector alone (Fig. S1).

**Fig. 2.**
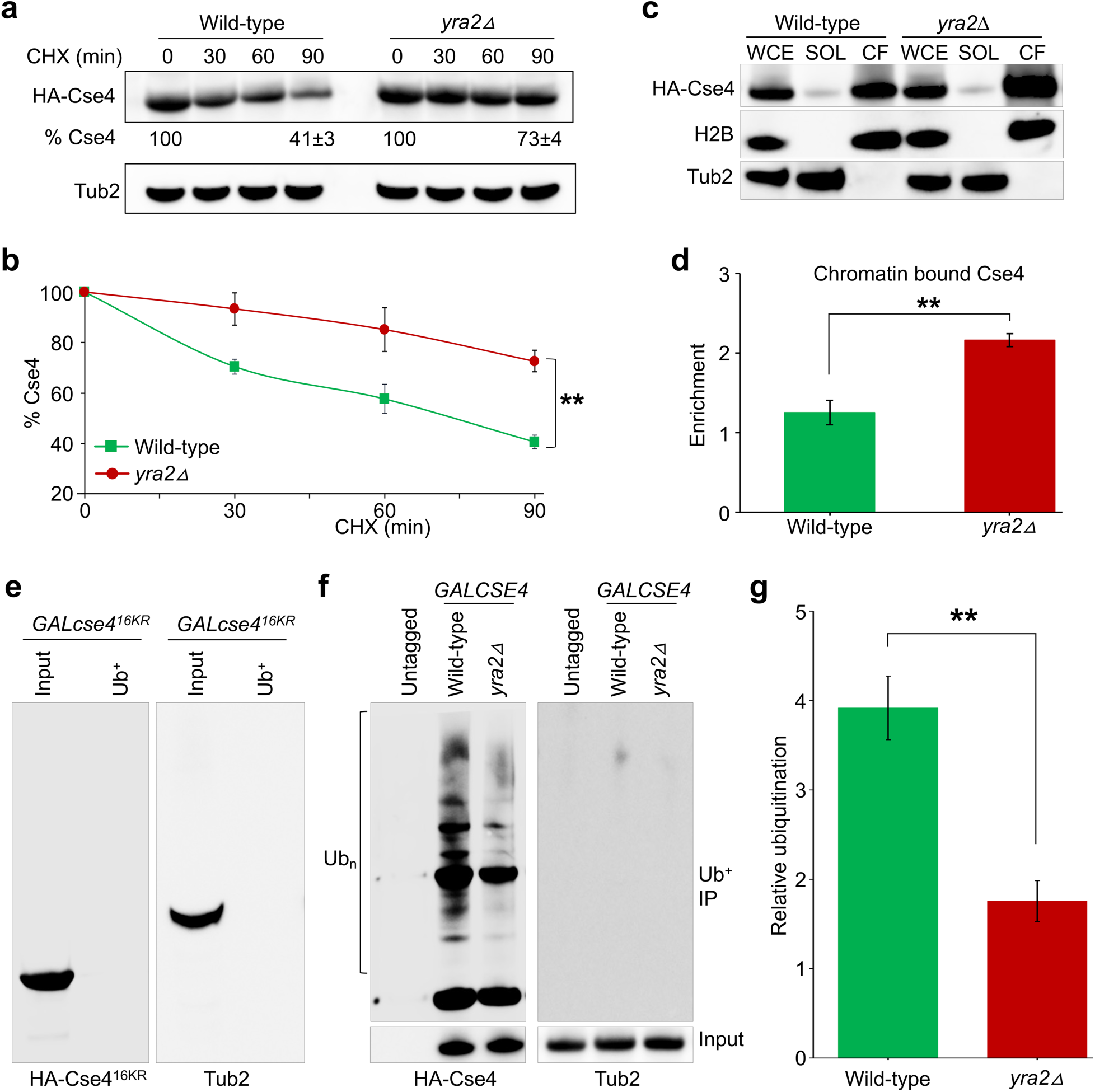
Deletion of *YRA2* results in increased protein stability and chromatin enrichment of Cse4 upon its overexpression. a) Increased stability of *GALCSE4* in *yra2Δ* strain. Western blot analysis was performed with whole cell extracts from wild-type (YMB12794) and *yra2Δ* (YMB12792) expressing *GALCSE4* (pMB1458) grown in galactose+raffinose (2% each) medium for 3 hours at 25°C and probed with anti-HA (HA-Cse4) and anti-Tub2 antibodies (loading control). Percentage of remaining Cse4 (normalized to Tub2) at the 90 minutes after CHX treatment derived from three biological repeats is shown as average± SE. b) Line graphs depicting percent Cse4 remaining at different time points after CHX treatment experiments from (a). Results from three biological experiments are shown as average±SE, \*\**p* value <0.01, Student’s t-test. c) Stabilized Cse4 is enriched in chromatin. Whole cell extracts (WCE), soluble (SOL) and chromatin fraction (CF) from wild-type (YMB12794), and *yra2Δ* (YMB12792) expressing *GALCSE4* (pMB1458) grown in galactose+raffinose (2% each) medium for 3 hours at 25°C were analyzed by western blotting using anti-HA (HA-Cse4), anti-Tub2, and anti-H2B antibodies. Tub2 and histone H2B were used as markers for soluble and chromatin fractions, respectively. d) Chromatin enrichment of Cse4 is significantly increased in *yra2Δ* strain. Enrichment as a ratio of Cse4 in chromatin fraction over input and normalized to H2B were calculated from intensity values calculated from western blots using Image J (SCHNEIDER *et al*. 2012). Average±SE from three biological replicates is shown. \*\**p* value <0.01, Student’s t-test. e) and f) Cse4 ubiquitination is reduced in *yra2Δ GALCSE4* strains. Western blots showing the levels of Cse4 ubiquitination (Ub_n_) in wild-type (YMB12794) and *yra2Δ* (YMB12792) strains carrying *GALCSE4* (pMB1458) after growth for 3 hours at 25°C in SC-Ura with galactose+raffinose (2% each). Wild-type strain with mutant *cse4^16KR^* in which all lysine residues are changed to arginine (YMB12815) and an untagged strain (YMB12793) were used as control in ubiquitin affinity pull down assays. Eluted proteins were analyzed by western blotting with anti-HA (Cse4) or anti-Tub2 (served as a loading control) antibodies. (g) Quantification of relative ubiquitination of Cse4. Ratio of ubiquitinated Cse4 (bracket in panel f) to the total Cse4 levels (input) in wild-type and *yra2Δ* strains is shown. The histogram represents the average from three biological replicates with SE. \*\**p* value <0.01, Student’s t-test.

To determine whether the higher stability, and chromatin enrichment of Cse4 are due to defects in ubiquitination, we assayed the levels of ubiquitinated Cse4 in wild-type and *yra2Δ* strain. Controls included a strain expressing *GALcse4^16KR^*, a mutant allele of Cse4 that cannot be ubiquitinated because all lysine residues are replaced with arginine (RANJITKAR *et al*. 2010; AU *et al*. 2013), and wild-type strain with untagged Cse4. Agarose beads with tandem ubiquitin-binding entities (Ub^+^) were used in an affinity assay (HJERPE *et al*. 2009; AU *et al*. 2013) to pull down ubiquitin-associated proteins and ubiquitination status of Cse4 was examined by western blot analysis. Control *GALcse4^16KR^*strain do not show laddering pattern for Cse4 (Fig. 2e). As reported previously (AU *et al*. 2013), ubiquitinated Cse4 is detected as a laddering pattern in wild-type strain expressing HA-Cse4, and no signal was observed in an untagged control strain (Fig. 2f). The laddering pattern representing ubiquitinated forms of Cse4 was reduced in the *yra2Δ* strain (Fig. 2f). Quantification of fraction of ubiquitinated Cse4 normalized to the total Cse4 levels (input) as described previously (AU *et al*. 2013) showed significantly reduced levels of ubiquitinated Cse4 in *yra2Δ* strain when compared to the wild-type strain (Fig. 2g). Taken together, these results support a role of Yra2 in ubiquitin-mediated proteolysis of Cse4.

### Yra2 facilitates the interaction of Cse4 with Psh1 for Cse4 proteolysis

The reduced ubiquitination of Cse4 in *yra2Δ* strain prompted us to examine whether Yra2 facilitates the interaction between Cse4 and E3 ubiquitin ligase Psh1, which is a major regulator of proteolysis of Cse4 (HEWAWASAM *et al*. 2010; RANJITKAR *et al*. 2010; MISHRA *et al*. 2015; CIFTCI-YILMAZ *et al*. 2018). IP experiments revealed that the *in vivo* interaction between Psh1-TAP and HA-Cse4 was reduced significantly in *yra2Δ* strains compared to that observed in wild-type strain (*p* value = 0.0093; Fig. 3a and b). We reasoned that overexpression of *PSH1* will suppress the SDL of *GALCSE4* and defects in Cse4 proteolysis in *yra2Δ* strains. Overexpression of *PSH1* suppressed the lethality of *GALCSE4* on galactose medium (Fig. 3c, *p* value = 0.002) and proteolysis of Cse4 proteolysis in *yra2Δ* strains (*p* value = 0.0079; Fig. 3d and e). Based on these results, we conclude that Psh1 mediated ubiquitination of Cse4 plays a role in SDL and Cse4 stability phenotypes observed in *yra2Δ GALCSE4* strains.

**Fig. 3.**
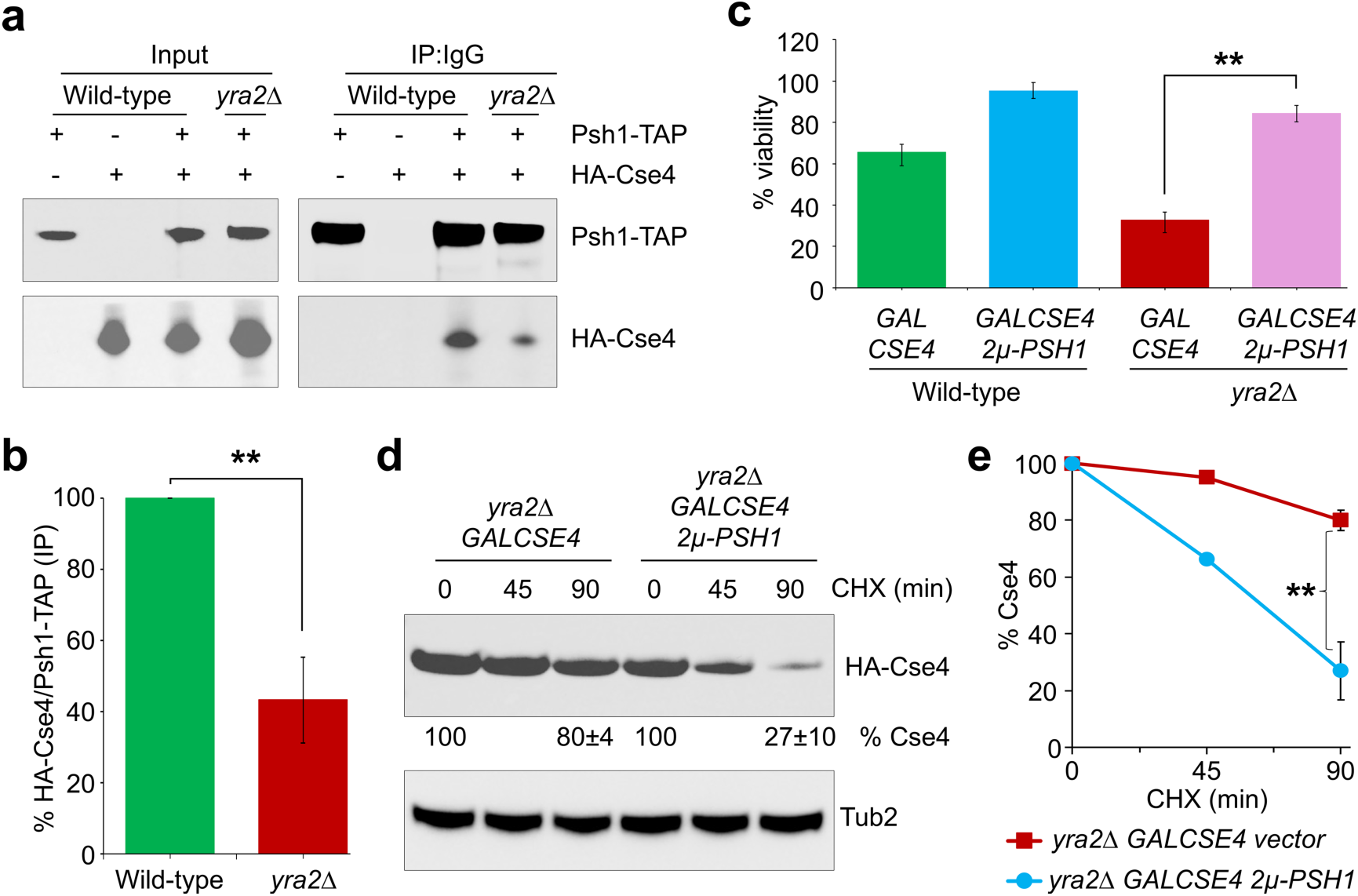
Yra2 facilitates Psh1-mediated ubiquitination and proteolysis of Cse4. a) *In vivo* interaction of Cse4 and Psh1 is reduced in *yra2Δ* strain. Wild-type with *GALCSE4* (YMB12794), Psh1-TAP with either vector (YMB10035) or *GALCSE4* (YMB10036) and Psh1-TAP *yra2Δ* strains with *GALCSE4* (YMB13039) were grown to the logarithmic phase and Cse4 expression induced by growth in SC-Ura with galactose+raffinose (2% each) medium for 3 hours at 25°C. IP experiments were performed using IgG agarose, and western blots were probed with anti-HA and anti-TAP (CAB1001, Thermo Scientific) antibodies. b) Quantitation show reduced interaction of Cse4 and Psh1 in *yra2Δ* strain. Western blots from (a) were quantified to determine the interaction between Psh1-TAP and HA-Cse4. The ratios showing levels of HA-Cse4 over Psh1-TAP from IP samples were calculated and normalized to a value of 100 for the wild-type strain. Error bars represent SE from three biological replicates. \*\**p* value <0.01, Student’s t-test. c) Overexpression of Psh1 suppresses the lethality caused by *GALCSE4* in *yra2Δ* strains. Viability assays were performed with wild-type with vector + *GALCSE4* (YMB13018) or *PSH1*/2*µ* + *GALCSE4* (YMB13019) and *yra2Δ* with vector + *GALCSE4* (YMB13020) or *PSH1*/2*µ* + *GALCSE4* (YMB13021). At least 500 cells from each strain were plated on glucose (2%) and galactose+raffinose (2% each) containing medium. Percent viability represents the ratio of colonies on galactose over glucose plates. Average±SE from three biological replicates is shown. \*\**p* value <0.01, Student’s t-test. d) Psh1 overexpression promotes proteolysis of Cse4 in *yra2Δ* strains. Western blotting was performed using whole cell extracts from *yra2Δ* with vector + *GALCSE4* (YMB13020) or *PSH1*/2*µ* + *GALCSE4* (YMB13021). Expression of Cse4 was induced in selective medium with galactose+raffinose (2% each) for 3 hours at 25°C and CHX treatment was performed as described in the *Materials and methods*. Western blots were probed with anti-HA (HA-Cse4) and anti-Tub2 antibodies (loading control). e) Line graphs depicting percent Cse4 remaining at different time points after CHX treatment. Western blots from (d) were quantified and results from three biological experiments are shown as average±SE, \*\**p* value <0.01, Student’s t-test.

### Yra2 prevents mislocalization of Cse4 to non-centromeric regions

Defects in Cse4 proteolysis contributes to *GALCSE4* SDL and this correlates with mislocalization of Cse4 to non-centromeric regions in *psh1Δ, slx5Δ, cdc48-3*, *met30-6*, *cdc4-1*, *cdc7-7*, *mck1Δ, rcy1Δ* or *hir2Δ* strains (HEWAWASAM *et al*. 2010; RANJITKAR *et al*. 2010; AU *et al*. 2013; CHENG *et al*. 2016; HILDEBRAND AND BIGGINS 2016; OHKUNI *et al*. 2016; CHENG *et al*. 2017; CIFTCI-YILMAZ *et al*. 2018; AU *et al*. 2020; EISENSTATT *et al*. 2020; OHKUNI *et al*. 2022; ZHANG *et al*. 2024). Hence, we performed ChIP-qPCR experiments to examine the localization of Cse4 in wild-type, and *yra2Δ* strains to control regions such as centromere (*CEN3)* and well characterized regions of Cse4 mislocalization such as promoters (*RDS1*, *SAP4*, and *SLP1*), coding region (*GUP2*) and non-coding intergenic region (*UGA3*) (OHKUNI *et al*. 2020; EISENSTATT *et al*. 2021). No significant differences in the levels of Cse4 were detected at *CEN3* in wild-type and *yra2Δ* strains (Fig. 4a). However, the enrichment of Cse4 at non-centromeric regions such as the promoters, coding and intergenic regions was significantly higher in the *yra2Δ* strain than the wild-type strain (Fig. 4a). We hypothesized that mislocalization of Cse4 to non-centromeric regions may contribute to reduced levels of histone H3 at these regions. In agreement with our hypothesis, ChIP-qPCR experiments showed reduced levels of histone H3 at non-centromeric regions of Cse4 mislocalization (Fig. 4b). No significant enrichment of histone H3 was detected at *CEN3* (Fig. 4b). We conclude that Yra2 prevents mislocalization of Cse4 to non-centromeric regions.

**Fig. 4.**
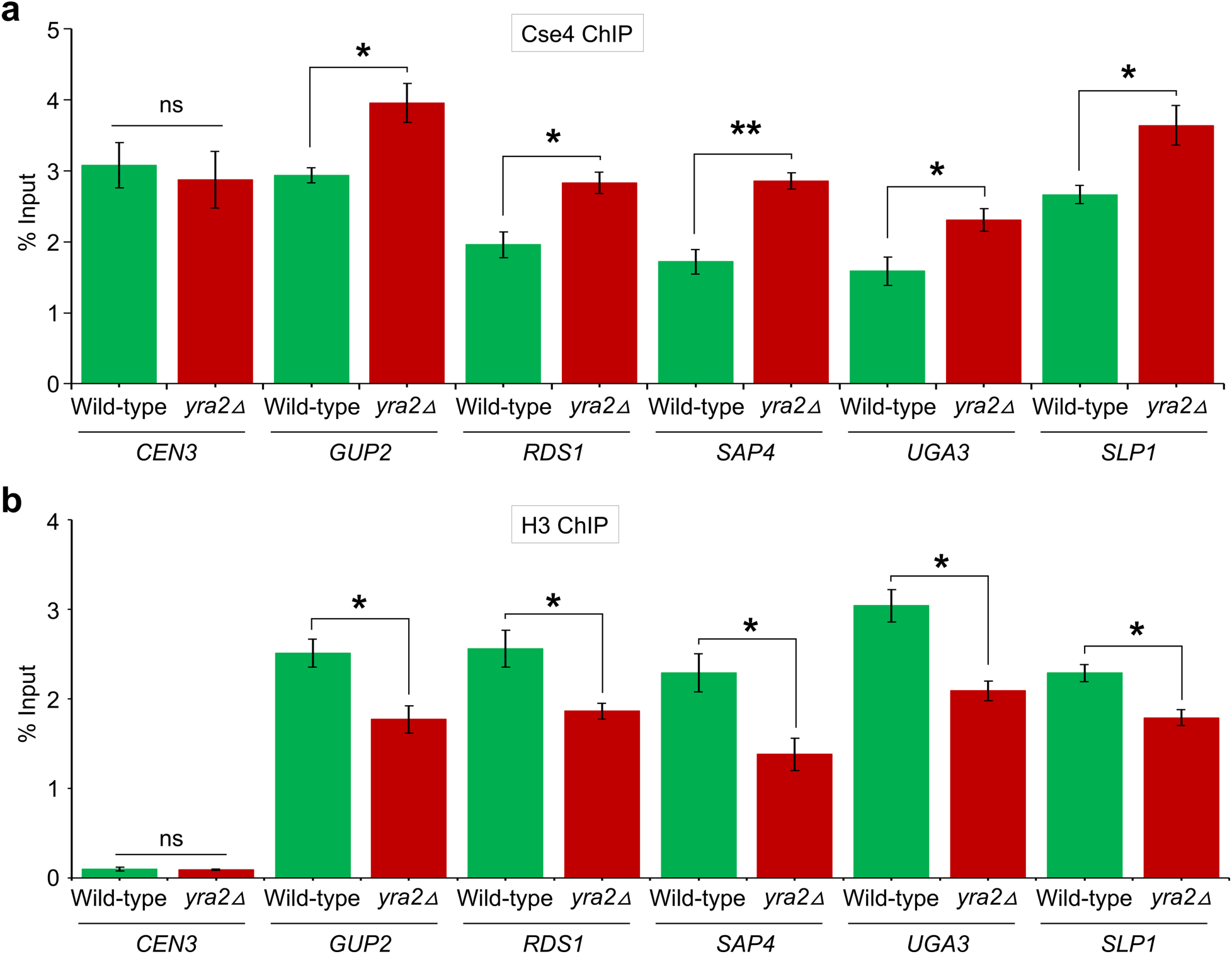
Cse4 is mislocalized to non-centromeric regions in *yra2Δ* strains. a) Levels of Cse4 are increased at non-centromeric regions in *yra2Δ GALCSE4* strains. ChIP experiments for HA-Cse4 and histone H3 were performed using chromatin prepared from wild-type (YMB12794) and *yra2Δ* (YMB12792) strains carrying *GALCSE4* (pMB1458) after growth for 3 hours at 25°C in SC-Ura with galactose+raffinose (2% each). Immunoprecipitation was performed using anti-HA agarose beads (for HA-Cse4) and anti-histone H3 antibodies as described in the *Materials and methods*. Enrichment of Cse4 at *CEN3* and non-centromeric *GUP2, RDS1, SAP4, UGA3,* and *SLP1* was determined by qPCR and is shown as % input. Average±SE from three biological replicates is shown. \**p* value <0.015, \*\**p* value <0.01, ns = statistically not significant, Student’s t-test. b) Levels of histone H3 are reduced at non-centromeric regions occupied by Cse4 in *yra2Δ GALCSE4* strains. Enrichment of histone H3 at *CEN3* and non-centromeric *GUP2, RDS1, SAP4, UGA3,* and *SLP1* was determined by qPCR and is shown as % input. Average±SE from three biological replicates is shown. \**p* value <0.015, ns = statistically not significant, Student’s t-test.

### Deletion of histone H4 alleles suppresses the SDL of *yra2Δ GALCSE4* strain

We have previously shown that deletion of either one of the histone H4 alleles (*HHF1* or *HHF2*) suppresses the SDL in a *psh1Δ GALCSE4* strain due to reduced interaction of Cse4 with histone H4 which contributes to reduced mislocalization of Cse4 (EISENSTATT *et al*. 2021). We examined the effect of dosage of histone H4 on the SDL phenotype of *yra2Δ GALCSE4* strain and Cse4-H4 interaction. *HHF1* and *HHF2* were deleted in wild-type and *yra2Δ* strains and growth phenotype was examined with and without *GALCSE4.* The SDL phenotype observed in *yra2Δ GALCSE4* strain was suppressed by deletion of either one of the H4 allele (Fig. 5a). To examine Cse4-H4 interactions, IP experiments were done with anti-HA agarose using whole cell extracts from wild-type, *yra2Δ, hhf2Δ* and *yra2Δ hhf2Δ* strains expressing *GAL1 HA-CSE4* (Fig. 5b). The relative levels of *in vivo* interactions between Cse4 and H4 was not significantly different between wild-type and *yra2Δ* strains, although slightly higher levels were observed in the *yra2Δ* strain (Fig. 5b and c). However, a significant reduction in the levels of interaction between Cse4 and H4 was observed in *hhf2Δ* and *yra2Δ hhf2Δ* strains when compared to the wild-type or *yra2Δ* strains (Fig. 5b and c). Importantly, the steady state levels of H4 and Cse4 were not affected in all the strains examined (Fig. S2). Based on these results, we conclude that the gene dosage of histone H4 dictates the extent of the Cse4-H4 interaction, which ultimately contributes to the accumulation of Cse4 in chromatin resulting SDL phenotype in *yra2Δ GALCSE4* strain.

**Fig. 5.**
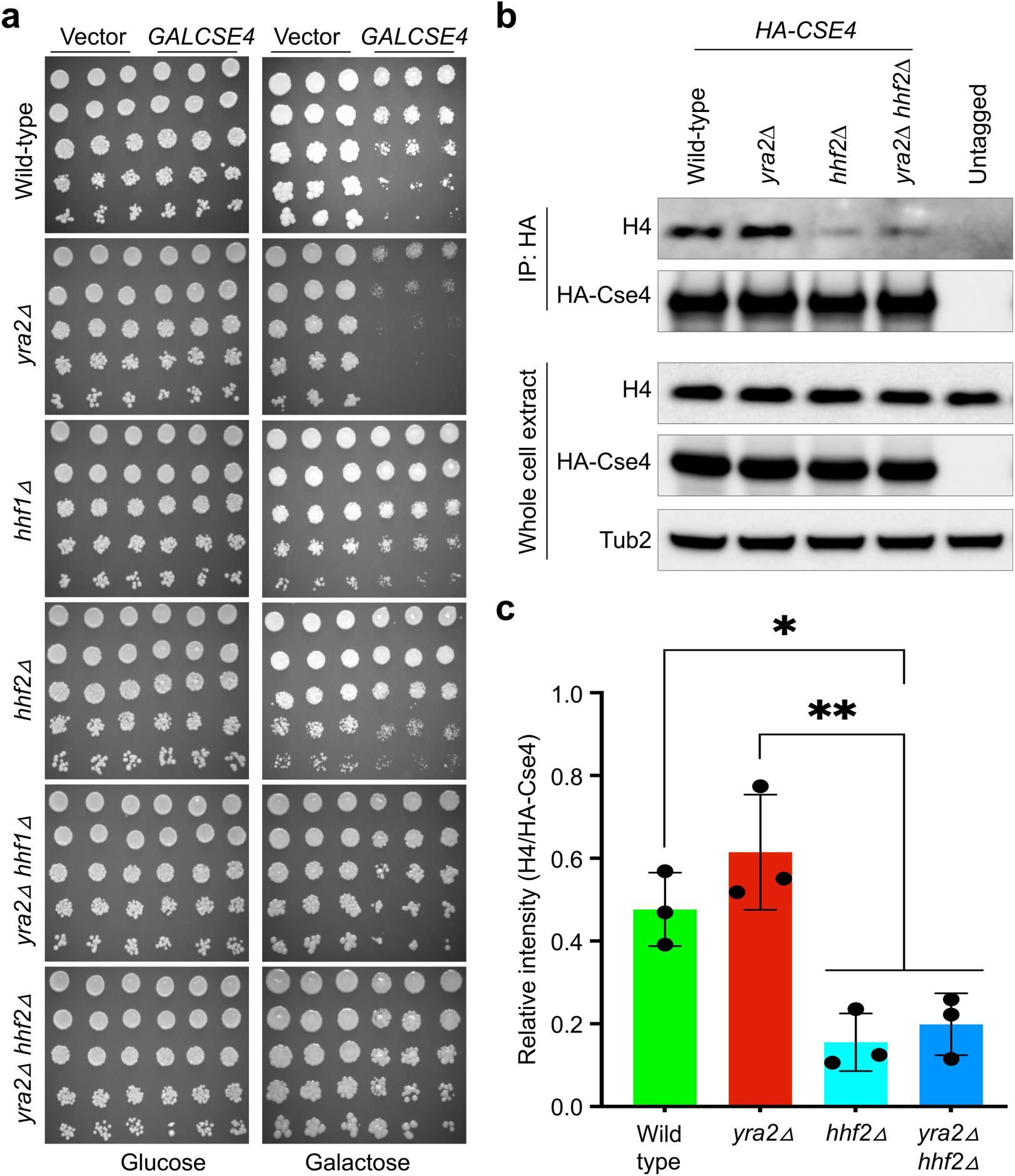
Dosage of histone H4 contributes to SDL in *yra2Δ GALCSE4* strains. a) *GALCSE4* induced SDL in *yra2Δ* strains is suppressed by deletion of either of histone H4 alleles. Wild-type with vector (YMB12793) or *GALCSE4* (YMB12794), *yra2Δ* with vector (YMB12791) or *GALCSE4* (YMB12792), *hhf1Δ* with vector (YMB12972) or *GALCSE4* (YMB12974), *hhf2Δ* with vector (YMB12973) or *GALCSE4* (YMB12975), *yra2Δ hhf1Δ* with vector (YMB12976) or *GALCSE4* (YMB12978), and *yra2Δ hhf2Δ* with vector (YMB12977) or *GALCSE4* (YMB12979) were grown to logarithmic phase, five-fold serial dilutions were prepared and plated on SC-Ura plates containing either glucose (2%) or galactose +raffinose (2% each) at 25°C. b) *In vivo* interaction of Cse4 and histone H4 is reduced in *yra2Δ* when compared to the wild-type strain. Wild-type (YMB12794), *yra2Δ* (YMB12792), *hhf2Δ* (YMB12975), and *yra2Δ hhf2Δ* (YMB12979) carrying *GALCSE4* (pMB1458) were grown to logarithmic phase, and Cse4 expression was induced by growth in SC-Ura with galactose+raffinose (2% each) medium for 4 hours at 25°C. IP experiments were performed using anti-HA agarose beads, and western blots were probed with anti-HA, histone H4 and anti-Tub2 antibodies. c) Quantitation show reduced interaction of Cse4 and H4 in *yra2Δ* strain. Western blots from (b) were quantified to determine the interaction between HA-Cse4 and H4. The ratios showing levels of H4 over HA-Cse4 from IP samples from three biological replicates are shown. \**p* value <0.05, \*\**p* value <0.01, Student’s t-test.

### RNA binding domain of Yra2 does not contribute to *GALCSE4 SDL* and Cse4 proteolysis

Our results show that *yra1-2* and *mex67-5* strains that exhibit defects in nuclear poly(A)^+^ RNA levels do not exhibit *GALCSE4* SDL strain, whereas *yra2Δ* exhibits *GALCSE4* SDL without defects in nuclear poly(A)^+^ RNA level. To further define the role of Yra2 in Cse4 proteolysis, we examined its primary sequence. Yra2 shares a conserved N-box and RNA binding domain with its family member Yra1 (ZENKLUSEN *et al*. 2001). However, Yra2 features a unique, evolutionarily conserved C-terminus chromatin target of PRMT1 (ChTOP) domain (amino acids 145-203) (https://www.yeastgenome.org/locus/S000001697/protein) (Fig. 6a), which is entirely absent in Yra1 (ZENKLUSEN *et al*. 2001; VAN DIJK *et al*. 2010; FANIS *et al*. 2012). We tested whether a mutant strain lacking this ChTOP domain, but an intact RNA binding domain (Fig. 6a) would mimic the *yra2*Δ phenotypes. As expected, no growth defects were observed on galactose medium for strains transformed with vector alone (Fig. 6b). Growth assays showed that *yra2-chtopΔ* strain exhibits *GALCSE4* SDL on galactose medium (Fig. 6b). To assess whether the ChTOP domain contributes to proteolysis of Cse4, we next examined stability of *GALCSE4* after induced expression for 3 hours in galactose-containing medium, followed by the addition of glucose and CHX. Western blot analysis revealed that Cse4 was rapidly degraded in the wild-type strain, whereas Cse4 stability was significantly increased in the *yra2-chtopΔ* strain (p < 0.01) (Fig. 6c and d). No significant difference in Cse4 stability was observed between the *yra2Δ* and *yra2-chtopΔ* strains (Fig. 6d). Based on these results, we conclude that the conserved RNA binding domain of Yra2 is dispensable for Cse4 proteolysis.

**Fig. 6.**
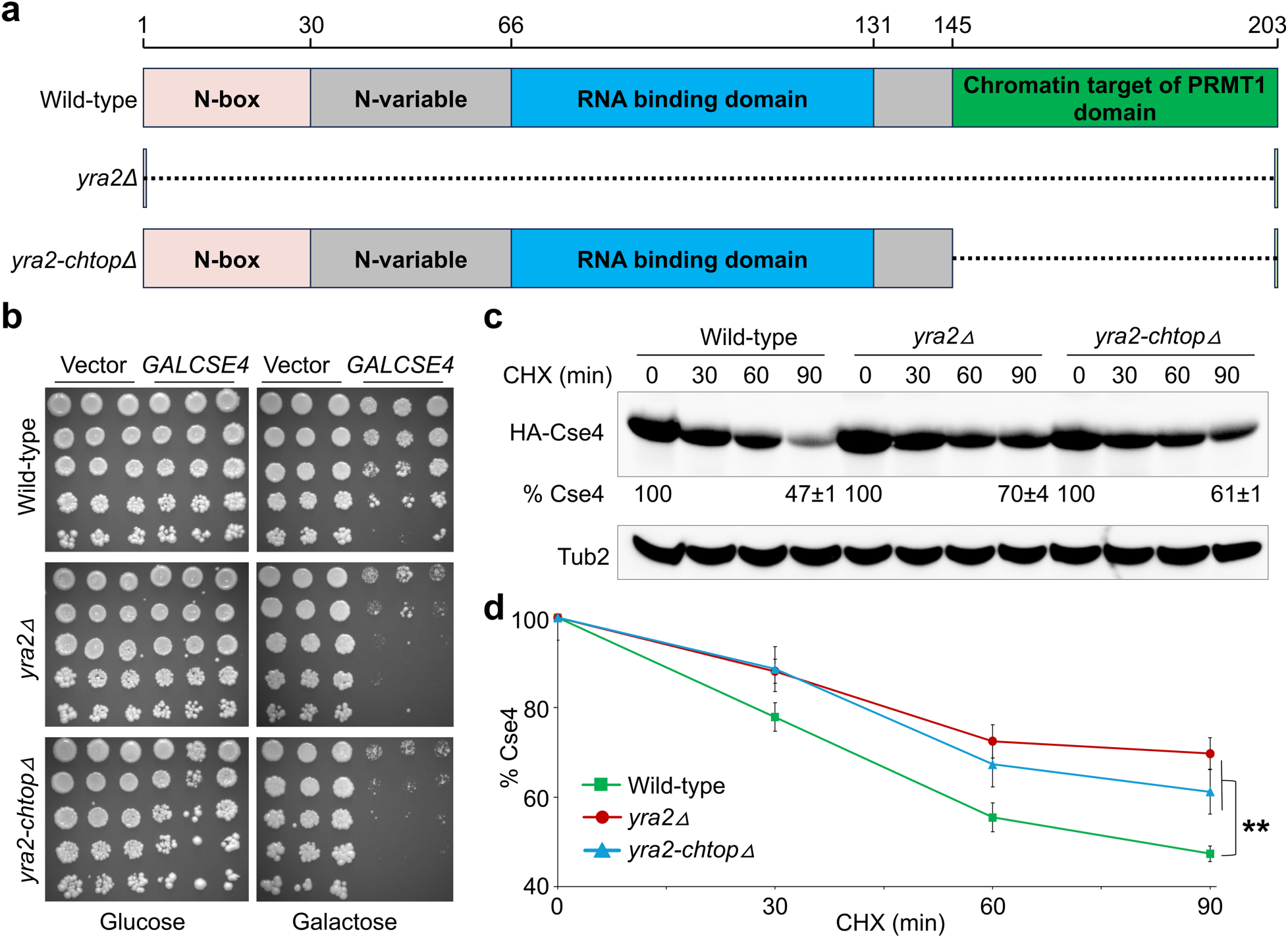
RNA binding domain of Yra2 does not contribute to *GALCSE4* SDL and Cse4 proteolysis. a) Schematic of full-length Yra2 and its alleles depicting different domains. b) *GALCSE4* causes growth defects in *yra2-chtopΔ* strain. Wild-type with vector (YMB12793) or *GALCSE4* (YMB12794), *yra2Δ* with vector (YMB12791) or *GALCSE4* (YMB12792), and *yra2-chtopΔ* with vector (YMB12951) or *GALCSE4* (YMB12952) were grown to logarithmic phase, five-fold serial dilutions were prepared and plated on SC-Ura plates containing either glucose (2%) or galactose+raffinose (2% each) at 25°C. c) Increased stability of *GALCSE4* in *yra2-chtopΔ* strain. Western blot analysis was performed with whole cell extracts from wild-type (YMB12794), *yra2Δ* (YMB12792), and *yra2-chtopΔ* (YMB12952) expressing *GALCSE4* (pMB1458) grown in galactose+raffinose (2% each) medium for 3 hours at 25°C and probed with anti-HA (HA-Cse4) and anti-Tub2 antibodies (loading control). d) Line graphs depicting percent Cse4 remaining at different time points after CHX treatment experiments from (c). Results from three biological experiments are shown as average±SE, \*\**p* value <0.01, Student’s t-test.

### Mislocalization of endogenous Cse4 contributes to defects in chromosome segregation in *yra2Δ* strain

Mislocalization of Cse4 and its homologs contributes to CIN in budding yeast, fission yeast, fruit flies and human cells (AU *et al*. 2008; SHRESTHA *et al*. 2017; CIFTCI-YILMAZ *et al*. 2018; ARISTIZABAL-CORRALES *et al*. 2019; MORENO-MORENO *et al*. 2019; AU *et al*. 2020; SHRESTHA *et al*. 2021; BALACHANDRA *et al*. 2024; SETHI *et al*. 2024; BALACHANDRA *et al*. 2025; SETHI *et al*. 2025). Our results for mislocalization of Cse4 in *yra2Δ* strains prompted to examine the role of Yra2 in chromosome segregation in the context of endogenous Cse4. We deleted *YRA2* in a strain carrying a reporter chromosome (RC) and quantified RC loss using a colony color assay, in which loss of the RC results in red sectors in otherwise white colonies (SPENCER *et al*. 1990). Colonies that were at least half red, indicative of RC loss during the first cell division, were scored. The frequency of RC loss in *yra2Δ* strains was approximately sixfold higher than in the wild-type strain (Fig. 7a). This increase in RC loss in *yra2Δ* strains is comparable to that reported for deletions of kinetochore genes (KASTENMAYER *et al*. 2005; MA *et al*. 2012). Our results for enrichment of overexpressed Cse4 in *yra2Δ* strain (Fig. 2c and d) prompted us to examine if endogenous Cse4 is enriched in *yra2Δ* strains. Our results showed that endogenous Cse4 was enriched in chromatin in the *yra2Δ* strain when compared to the wild-type strain (Fig. S3a and b). Chromosome segregation defects due to mislocalization of endogenous Cse4 are suppressed by constitutive expression of histone H3 (*Δ16H3*) (AU *et al*. 2008). We determined that frequency of RC loss in the *yra2Δ* strain was significantly reduced upon expression of *Δ16H3* (*p* value = 0.0051, Fig. 7b). The frequency of RC loss in *yra2Δ Δ16H3* strain was similar to that observed for the wild-type strain (Fig. 7b). Taken together, we conclude that mislocalization of endogenous Cse4 contributes to defects in chromosome segregation in *yra2Δ* strain.

**Fig. 7.**
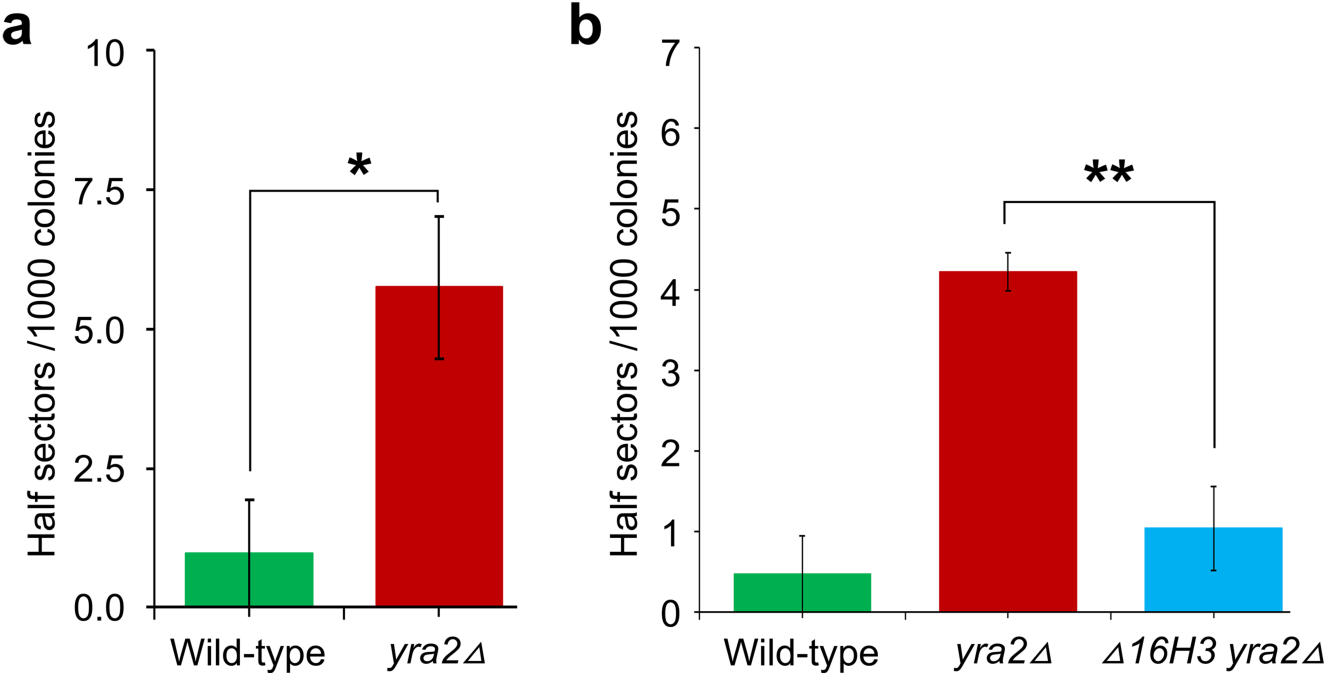
Yra2 is required for faithful chromosome segregation. a) Deletion of *YRA2* results in a CIN phenotype. Frequency of CIN in wild-type (YPH1018), and *yra2Δ* (YMB13065) strains was determined as described in the *Materials and methods*. About 1000 colonies were counted and average±SE from three transformants is shown. \**p* value <0.05, Student’s t-test. b) Increased CIN in *yra2Δ* is suppressed by constitutive expression of histone H3 (*Δ16H3*). Frequency of CIN in wild-type (YPH1018), *yra2Δ* (YMB13065), and *yra2Δ Δ16H3* (YMB12998) strains was determined as described in the *Materials and methods*. About 1000 colonies were counted and average±SE from three transformants is shown. \*\**p* value <0.01, Student’s t-test.

## Discussion

The evolutionarily conserved histone H3 variant Cse4 and its homologs play a critical role in ensuring faithful chromosome segregation across eukaryotic organisms (KITAGAWA AND HIETER 2001; MERALDI *et al*. 2006; VERDAASDONK AND BLOOM 2011; BURRACK AND BERMAN 2012; MADDOX *et al*. 2012; BIGGINS 2013; BLOOM AND COSTANZO 2017; FRIEDMAN AND FREITAG 2017; BARRA AND FACHINETTI 2018; SIDHWANI AND STRAIGHT 2023; SALINAS-LUYPAERT AND FACHINETTI 2024; DUTTA *et al*. 2025; GRECO *et al*. 2026). Genome wide genetic screens have uncovered a role for ubiquitin ligases, protein kinases, and histone chaperones in preventing mislocalization of Cse4 (HEWAWASAM *et al*. 2010; RANJITKAR *et al*. 2010; CIFTCI-YILMAZ *et al*. 2018; AU *et al*. 2020; EISENSTATT *et al*. 2020; OHKUNI *et al*. 2022). In this study, we define a novel role for Yra2 in Cse4 proteolysis and preventing its mislocalization to non-centromeric regions, thereby preserving chromosomal stability. These conclusions are based on results showing that *yra2Δ GALCSE4* strain exhibits: a) growth defects, b) increased Cse4 stability, c) enrichment of Cse4 in chromatin, d) mislocalization of Cse4 to non-centromeric regions, and d) reduced ubiquitination of Cse4. Mechanistically, we have determined that Yra2 regulates an *in vivo* interaction of Cse4 with Psh1 for Cse4 proteolysis. Cse4 mislocalization and CIN phenotypes of *yra2Δ* strain are regulated by a dynamic balance of Cse4 and canonical histones H3 and H4 that prevent and promote mislocalization of Cse4, respectively. Several experimental evidence show that the role of Yra2 in Cse4 proteolysis are independent of its RNA-related functions.

Psh1 is one of the major E3 ubiquitin ligases that regulated proteolysis of overexpressed Cse4 (HEWAWASAM *et al*. 2010; RANJITKAR *et al*. 2010; CIFTCI-YILMAZ *et al*. 2018). The phenotypes of *yra2Δ GALCSE4* strain are similar to those observed for *psh1Δ GALCSE4* strain such as defects in Cse4 ubiquitination that contribute to stability of Cse4 with enrichment in chromatin and mislocalization to non-centromeric regions (HEWAWASAM *et al*. 2010; RANJITKAR *et al*. 2010; HEWAWASAM *et al*. 2014; HILDEBRAND AND BIGGINS 2016; CIFTCI-YILMAZ *et al*. 2018; EISENSTATT *et al*. 2020). Mechanistically, we have shown that Yra2 facilitates Cse4-Psh1 interaction and consistent with this observation, overexpression of *PSH1* suppresses *yra2Δ GALCSE4* SDL and Cse4 protein stability. Given the essential role for Cse4 in *CEN* function and cell viability, it is not surprising that cellular levels of Cse4 are fine-tuned by modulating the activity of a major ubiquitin ligase such as Psh1 by multiple factors such as HIR complex (CIFTCI-YILMAZ *et al*. 2018), Cdc7 kinase (EISENSTATT *et al*. 2020), FACT complex (DEYTER AND BIGGINS 2014), and Yra2 as discussed in this study.

Previous studies have established a role for stoichiometric balance among Cse4, H3 and H4 in preventing and promoting the mislocalization of Cse4 in budding yeast (AU *et al*. 2008; DEYTER *et al*. 2017; AU *et al*. 2020; EISENSTATT *et al*. 2021). For example, reduced gene dosage of histone H4 suppresses *GALCSE4* mediated SDL phenotype in *psh1Δ*, *slx5Δ*, *cdc4-1*, *doa1Δ*, *hir2Δ*, and *cdc7-4*, and *cdc48-3* strains (EISENSTATT *et al*. 2021; OHKUNI *et al*. 2022). Consistent with these studies, we observed suppression of SDL phenotype in *yra2Δ GALCSE4* strain upon deletion of either one of the histone H4 alleles. Furthermore, we have shown that CIN phenotype of *yra2Δ* strain are suppressed by constitutive overexpression of histone H3 (*Δ16H3*) (AU *et al*. 2008; AU *et al*. 2020). As described for other regulators of Cse4 proteolysis, we provide evidence showing that that a proper stoichiometric balance between Cse4 and canonical histones H3 and H4 are essential for cell survival and faithful chromosome segregation (EISENSTATT *et al*. 2021; OHKUNI *et al*. 2024).

Yra2 derived its name based on its interaction with RNA (ZENKLUSEN *et al*. 2001) and genetic interactions have been reported between *YRA1* and *YRA2* (ZENKLUSEN *et al*. 2001; ZENKLUSEN *et al*. 2002; KASHYAP *et al*. 2005). In this study, we show that the role for Yra2 in Cse4 proteolysis is independent of its RNA-related functions. Several experimental evidence support this conclusion. For example, unlike *yra1-2* and *mex67-5* mutants which show significantly elevated levels of nuclear poly(A)^+^ RNA foci, *yra2Δ* strain did not exhibit increased nuclear poly(A)^+^ RNA foci and *yra1-2 GALCSE4* and *mex67-5 GALCSE4* strains do not exhibit SDL. Notably, a recent study based on stoichiometric measurements of molecules present in messenger ribonucleoprotein particles suggest distinct functions for Yra1 and Yra2 in mRNA biogenesis and export (ASADA *et al*. 2023). Furthermore, *yra2-chtopΔ* strain with an intact RNA binding domain exhibits *GALCSE4* SDL and defects in Cse4 proteolysis. Notably, Yra2 is evolutionarily conserved and the human homolog ChTOP interacts with TREX complex components such as Alyref (Yra1 in yeast) and Uap56 (Mex67 in yeast) (CHANG *et al*. 2013). RNAi-mediated depletion of human ChTOP also does not cause significant nuclear accumulation of poly(A)^+^ RNA (CHANG *et al*. 2013), a phenotype consistent with our observations for poly(A)^+^ RNA in the *yra2Δ* strain. Based on these results, we conclude that the conserved RNA binding domain of Yra2 is dispensable for Cse4 proteolysis.

Our results showed that deletion of *YRA2* results in CIN phenotype that correlated with the increased chromatin enrichment of endogenous Cse4. We propose that the CIN is most likely linked to Cse4 mislocalization in *yra2Δ* strains because CIN phenotype was suppressed by constitutive expression of H3 (*Δ16H3*). Previous studies have shown that *Δ16H3* suppresses mislocalization and chromosome loss in mutants defective in proteolysis of endogenous Cse4, such as *met30-6*, *cdc4-1* and *cse4^16KR^*(AU *et al*. 2008; AU *et al*. 2020). Similarly, overexpression of histone H3 suppresses chromosome segregation defects caused by mislocalization of Cnp1 in fission yeast (KITAGAWA *et al*. 2014). Moreover, Yra2 interacts *in vivo* with the essential kinetochore protein Dsn1 (AKIYOSHI *et al*. 2010) and deletion of *YRA2* exhibits negative genetic interactions and growth defects when combined with mutants of kinetochore genes Dad2 (*dad2-9*), Mif2 (*mif2-3*), and Nnf1 (*nnf1-4*8) (COSTANZO *et al*. 2016), further supporting the role of Yra2 in chromosome segregation due to enhanced enrichment of Cse4 in chromatin.

In summary, our study uncovers a novel, RNA export-independent role for Yra2 in maintaining Cse4 homeostasis in budding yeast. Deletion of *YRA2* resulted in increased stability and chromatin enrichment of Cse4, reduced Cse4 ubiquitination, mislocalization of Cse4, and CIN. These defects are attributable to defects in the interaction between Cse4 and Psh1 as overexpression of *PSH1* suppresses SDL and Cse4 stability in *yra2Δ GALCSE4* strains. Overall, our findings establish Yra2 as a key factor preventing Cse4 mislocalization by promoting its ubiquitin-mediated degradation, thereby ensuring faithful chromosome segregation. These studies are clinically significant because overexpression of CENP-A is observed in various human cancers and this correlates with poor prognosis (TOMONAGA *et al*. 2003; AMATO *et al*. 2009; LI *et al*. 2011; MCGOVERN *et al*. 2012; SUN *et al*. 2016; ZHANG *et al*. 2016; XU *et al*. 2020; LI *et al*. 2022). Mislocalization of overexpressed CENP-A to non-centromeric regions has been reported in various cell lines (LACOSTE *et al*. 2014; ATHWAL *et al*. 2015; SHRESTHA *et al*. 2017). Induction of CENP-A overexpression alone, in defined cell lines and mouse model studies is sufficient to promote CIN (SHRESTHA *et al*. 2017; SHRESTHA *et al*. 2021), increase proliferation (SAHA *et al*. 2020), invasion (SHRESTHA *et al*. 2021), and tumor growth in mouse models (LING *et al*. 2024). Defining pathways that regulate cellular levels of Cse4 advance our understanding of how defects in proteolysis of CENP-A may contribute to aneuploidy in human cancers.

## Acknowledgements

We thank the members of Basrai laboratory for helpful discussion and comments on the manuscript. This research was supported by the Intramural Research Program (IRP) of the National Institutes of Health (NIH) to MAB (IRP project# ZIA BC 010822). The contributions of the NIH authors were made as part of their official duties as NIH federal employees, are in compliance with agency policy requirements, and are considered Works of the United States Government. However, the findings and conclusions presented in this paper are those of the authors and do not necessarily reflect the views of the NIH or the U.S. Department of Health and Human Services. M.C. and C.B. were supported by the NIH grant (RO1 HG005853), and Canadian Institutes of Health Research grant (PJT-180285). D.Z. was supported by Canadian Institutes of Health Research grant (PJT-192036).

## Abbreviations used

*CEN*: centromere
CENP-A: centromere protein-A
ChIP: chromatin immunoprecipitation
CIN: chromosomal instability
DDK: Dbf1-dependent kinase
IP: immunoprecipitation
PTMs: post-translational modifications
qPCR: quantitative PCR
SDL: synthetic dosage lethality
SGA: synthetic genetic array

## Supplemental Figures

**Fig. S1.**
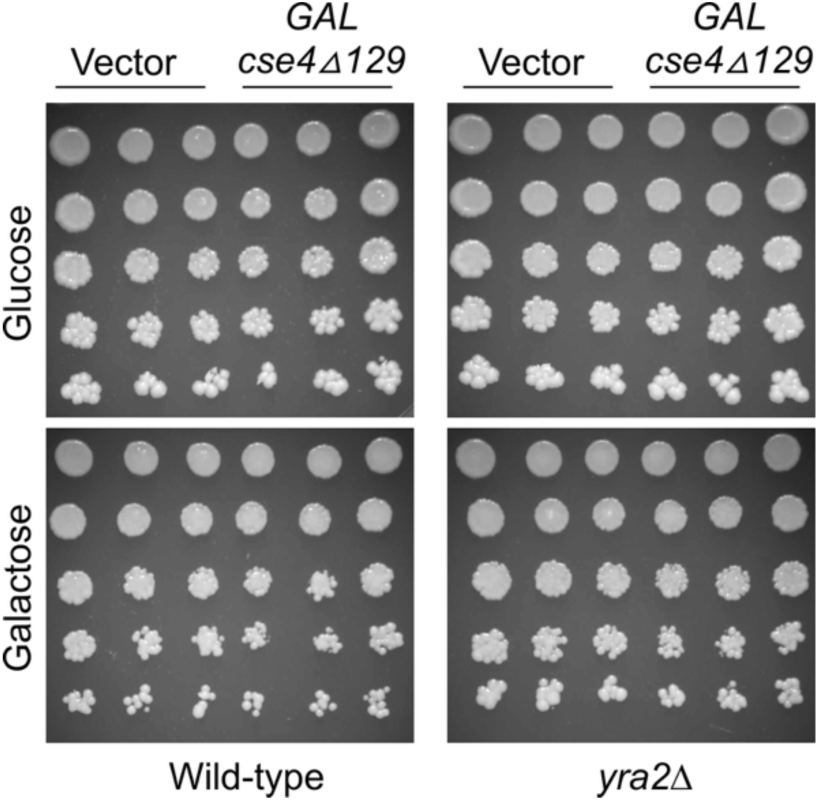
The N-terminus of Cse4 contributes to the *GALCSE4* induced SDL in *yra2Δ* strains. Wild-type with vector (YMB12793) or *GALcse4Δ129* (YMB12924) and the isogenic *yra2Δ* with vector (YMB12791) or *GALcse4Δ129* (YMB12927) were grown to logarithmic phase, five-fold serial dilutions were prepared and plated on SC-Ura plates containing either glucose (2%) or galactose+raffinose (2% each) at 25°C.

**Fig. S2.**
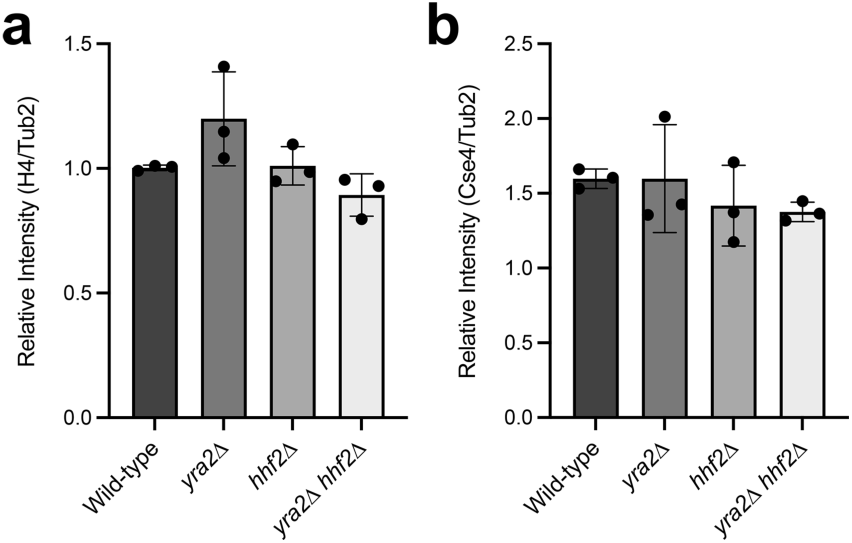
The steady state levels of histone H4 and Cse4 are not affected across different strains examined in this study (Related to Fig. 5).

**Fig. S3.**
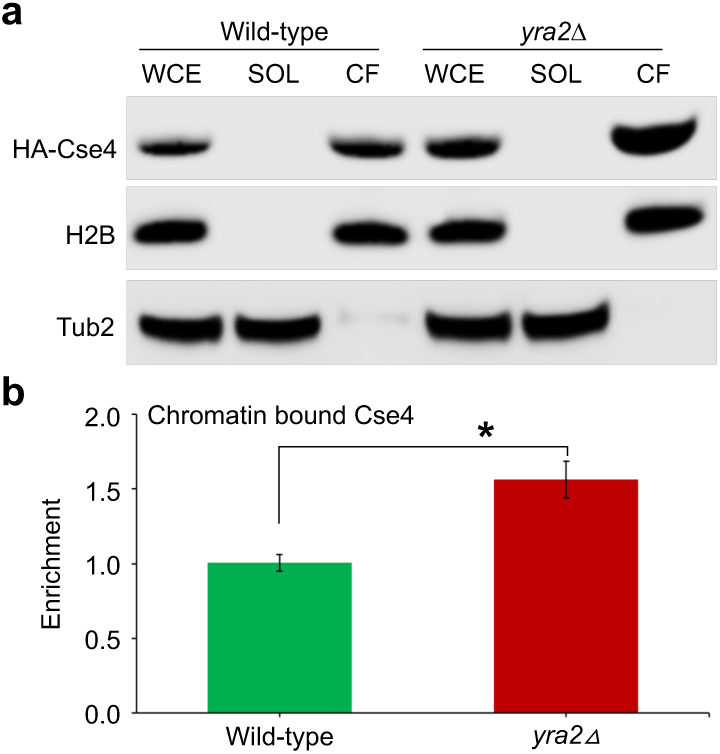
Endogenous Cse4 is enriched in chromatin in *yra2Δ* strains. a) Levels of chromatin bound endogenously expressed Cse4 in *yra2Δ* strain. Whole cell extracts (WCE), soluble (SOL) and chromatin fraction (CF) from wild-type (YMB10574), and *yra2Δ* (YMB13044) expressing HA-Cse4 from its native promoter at the endogenous locus were analyzed by western blotting using anti-HA (HA-Cse4), anti-Tub2, and anti-H2B antibodies. Tub2 and histone H2B were used as markers for soluble and chromatin fractions, respectively. b) Chromatin enrichment of Cse4 is significantly increased in *yra2Δ* strain. Enrichment as a ratio of Cse4 in chromatin fraction over input and normalized to H2B were determined from intensity values calculated from western blots using Image J (SCHNEIDER *et al*. 2012). Average±SE from three biological replicates is shown. \**p* value <0.05, Student’s t-test.

